# Single-Cell Transcriptomic Analysis Highlights Specific Cell Types of Wheat Manipulated by *Fusarium graminearum* Leading to Susceptibility

**DOI:** 10.1101/2024.06.08.598051

**Authors:** Wan-Qian Wei, Shuang Li, Dong Zhang, Wei-Hua Tang

## Abstract

Phytopathogenic fungi can be recognized by the plant immune system and trigger host defenses, but adapted pathogens cause susceptibility. How different cell types cooperate and orchestrate biological processes in response to heterogeneous colonization of organs by adapted and non-adapted pathogens remains largely unknown. Here we employed single-cell RNA sequencing to dissect the responses of wheat coleoptiles to infection by the adapted fungal pathogen *Fusarium graminearum* (*Fgr*) and the non-adapted fungal pathogen *Fusarium oxysporum* f. sp. *cubense* (*Foc*) at 1-, 2-, and 3-days post-inoculation. We profiled the transcriptomes of over 90,000 cells and identified eight major cell types in coleoptiles: stomata, epidermis, chlorenchyma, parenchyma, outer sheath, inner sheath, phloem, and procambium. Differential expression analyses showed that the capacity of different cell types to respond to fungal infection varied. The upregulation of immune pathways was compartmentalized in nonhost resistance to *Foc*, but widespread in susceptible interaction with *Fgr*. Pseudotime analyses revealed continuous cell state transitions in the disease progression of infected cell types. Our work indicates that the phloem and outer sheath are specific cell types that collaborate for the rapid onset of nonhost resistance. *Fgr* induces a state of low transcriptional activity in the chlorenchyma. Cell trajectory analysis suggests that the competition between immune and susceptible processes in parenchyma results in specific cell states that are favored by the adapted pathogen *Fgr*. Overall, this work explains how cell types collaborate and are manipulated during fungal infections, providing insight into the intercellular mechanisms of plant immunity.

## Introduction

Plants have evolved a two-tiered innate immune system during their constant exposure to various microbes. In the first tier, pattern recognition receptors (PRRs) on the cell surface perceive conserved pathogen- or microbe-associated molecular patterns (PAMPs/MAMPs), leading to pattern-triggered immunity (PTI) (Jones and Dangl, 2006; Ngou et al., 2022). In the second tier, intracellular nucleotide-binding leucine-rich repeat receptors (NLRs) recognize pathogen effectors directly or indirectly, initiating effector-triggered immunity (ETI) (Jones and Dangl, 2006; Ngou et al., 2022). The activation of immune receptors ultimately leads to various downstream immune responses, such as Ca^2+^ influx, reactive oxygen species generation, MAP kinase activation, and production of phytohormones, thereby inducing massive transcriptional reprogramming (Ngou et al., 2022). Moreover, local PTI and ETI responses can further induce systemic acquired resistance (SAR) in distal tissues (Peng et al., 2021). Plants defend against numerous non-adapted pathogens through diverse mechanisms, including incompatible physicochemical traits, preformed chemical barriers, PTI, and ETI, collectively termed nonhost resistance (Ayliffe and Sørensen, 2019; Wu et al., 2023). However, adapted pathogens can evade host detection and suppress these immune responses by producing various effector proteins or metabolites, which leads to susceptibility (Jones and Dangl, 2006; Ngou et al., 2022). The understanding of molecular interactions between the plant immune system and pathogen effectors is increasing, but knowledge of how immune responses are allocated and coordinated in different plant cells is still limited.

Transcriptional responses of host plants reflect not only the induction of downstream defense pathways but also the activation of immune recognition and signaling pathways. This is because immune pathways are often self-amplified in transcriptional reprogramming (Li et al., 2016). Generally, pathogen distribution within host tissues is uneven, with multiple infection stages observed simultaneously, resulting in unequal symptom development (Zhu et al., 2023b). To address this, high-throughput single-cell RNA sequencing (scRNA-seq) was recently used to dissect plant responses to pathogens, revealing cellular heterogeneity in Arabidopsis-microbe interactions (Nobori et al., 2023; Zhu et al., 2023a; Delannoy et al., 2023; Tang et al., 2023). However, in crop plants, the spatial and temporal partitioning of various immune pathways and pathogen-caused susceptibility pathways among cells is still poorly understood.

Wheat (*Triticum aestivum*) is a staple crop that accounts for approximately 20% of the human caloric intake worldwide (FAO, 2019). However, wheat yields are significantly impacted by pathogens and pests, with annual losses estimated at 21.5% (Savary et al., 2019). Notably, Fusarium head blight (FHB) alone contributes to approximately 2.85% of these losses, ranking second after leaf rust in terms of impact (Savary et al., 2019). Despite the identification of numerous quantitative trait loci (QTL) offering varying degrees of FHB resistance, their effects are generally moderate or unstable (Gorash et al., 2021; Ma et al., 2022; Hu et al., 2022). In the absence of wheat varieties completely immune to FHB (Dweba et al., 2017; Hu et al., 2022; Moonjely et al., 2023), strategies focusing on modifying susceptibility genes present a viable alternative for FHB control. While these strategies to enhance resistance are well-documented in numerous plant-pathogen interactions (Garcia-Ruiz et al., 2021), research in the context of FHB remains limited (Fabre et al., 2020; Gorash et al., 2021).

*Fusarium graminearum* (*Fgr*), the primary causal agent of FHB, also induces head blight in barley, stalk rot in maize, and root rot in various crops, including wheat, maize, and soybean (Moonjely et al., 2023). The unequal symptom development in the wheat-*Fgr* interaction was also observed (Brown et al., 2010; Zhang et al., 2012; Qiu et al., 2019; Mentges et al., 2020). Transcriptomic studies of this interaction have primarily focused on elucidating disease resistance mechanisms conferred by QTLs in wheat (Xiao et al., 2013; Long et al., 2015; Dhokane et al., 2016; Biselli et al., 2018; Wang et al., 2018; Sari et al., 2019), with some attempting to identify susceptibility genes (Erayman et al., 2015; Chetouhi et al., 2016; Pan et al., 2018; Su et al., 2021). However, these analyses, conducted on bulk tissues, suffer from inherent biases, as averaging signals from individual cells can obscure key information. This bias, akin to Simpson’s Paradox in statistics, can result in qualitatively incorrect interpretations caused due to the failure to compartmentalize data by cell type (Trapnell, 2015). Asynchronous infection and uneven effector targeting result in heterogeneous cellular responses that bulk RNA-seq analyses cannot reveal, especially considering the complex tissue structure of wheat spikes. Hence, it is still uncertain how *F. graminearum* breaks down the immune responses coordinated by different cell types in wheat.

In this study, we profiled single-cell transcriptomes of wheat seedling coleoptiles in response to fungal pathogens. To elucidate how different cell types of wheat effectively orchestrate biological processes in nonhost resistance and identify specific cell types manipulated in susceptible interaction with *Fgr*, a non-adapted *Fusarium* pathogen was used as a positive control. Through single-cell transcriptomic analysis, we have elucidated that the upregulation of PTI, ETI, SAR, and secondary metabolism engages multiple cell types, illustrating the compartmentalization of the plant nonhost resistance. Among the coleoptile cell types, the phloem and outer sheath exhibit profound and distinct responses to pathogens during the rapid onset of nonhost resistance. However, the adapted fungal pathogens *Fgr* manipulated the chlorenchyma and parenchyma at different stages to facilitate colonization. In summary, our study elucidates how different cell types work together during nonhost resistance and how *Fgr* disrupts immune responses by manipulating specific cell types. This work also offers a foundational resource for identifying novel candidate genes in targeted cell types to develop FHB-resistant wheat cultivars while minimizing growth penalties.

## Results

### Adapted and non-adapted fungal pathogens exhibited comparable invasion patterns in the early stages, but with different tissue preferences

To elucidate the cellular responses of wheat to *Fusarium* infection, we used two fungal pathogens in the wheat-*Fusarium* pathosystem of the coleoptile, a leaf-like organ that encloses foliage leaves (Zhang et al., 2012). These pathogens include *Fusarium graminearum* (*Fgr*), a known wheat pathogen, and *Fusarium oxysporum* f. sp. *cubense* (*Foc*), a banana pathogen that is not naturally pathogenic to wheat (Zhang et al., 2019). The ancestral lineages of *Foc* and *Fgr* diverged approximately 35 million years ago (O’Donnell et al., 2013). These two *Fusarium* species share more than two-thirds of conserved genes (approximately 9000 conserved genes) and their orthologue genes have 85% nucleotide sequence identity on average (Ma et al., 2010), but *Foc* lacks some effectors such as fusaoctaxin which *Fgr* has (Jia et al., 2019). The adapted pathogen *Fgr* rapidly colonized the coleoptile within three to four days post-inoculation (DPI), whereas the non-adapted pathogen *Foc* only induced limited browning lesions upon inoculation of its conidia on wounded sites (Figure 1A). When supplemented with the effector non-ribosomal peptide fusaoctaxin from *Fgr*, the non-adapted pathogen *Foc* can cause disease on coleoptiles similar to that caused by *Fgr* (Jia et al., 2019). These findings suggest a notable overlap in pathogenic mechanisms between the two *Fusarium* species, making *Foc* a suitable positive control for elucidating effective defense responses of the wheat coleoptile.

**Figure 1.**
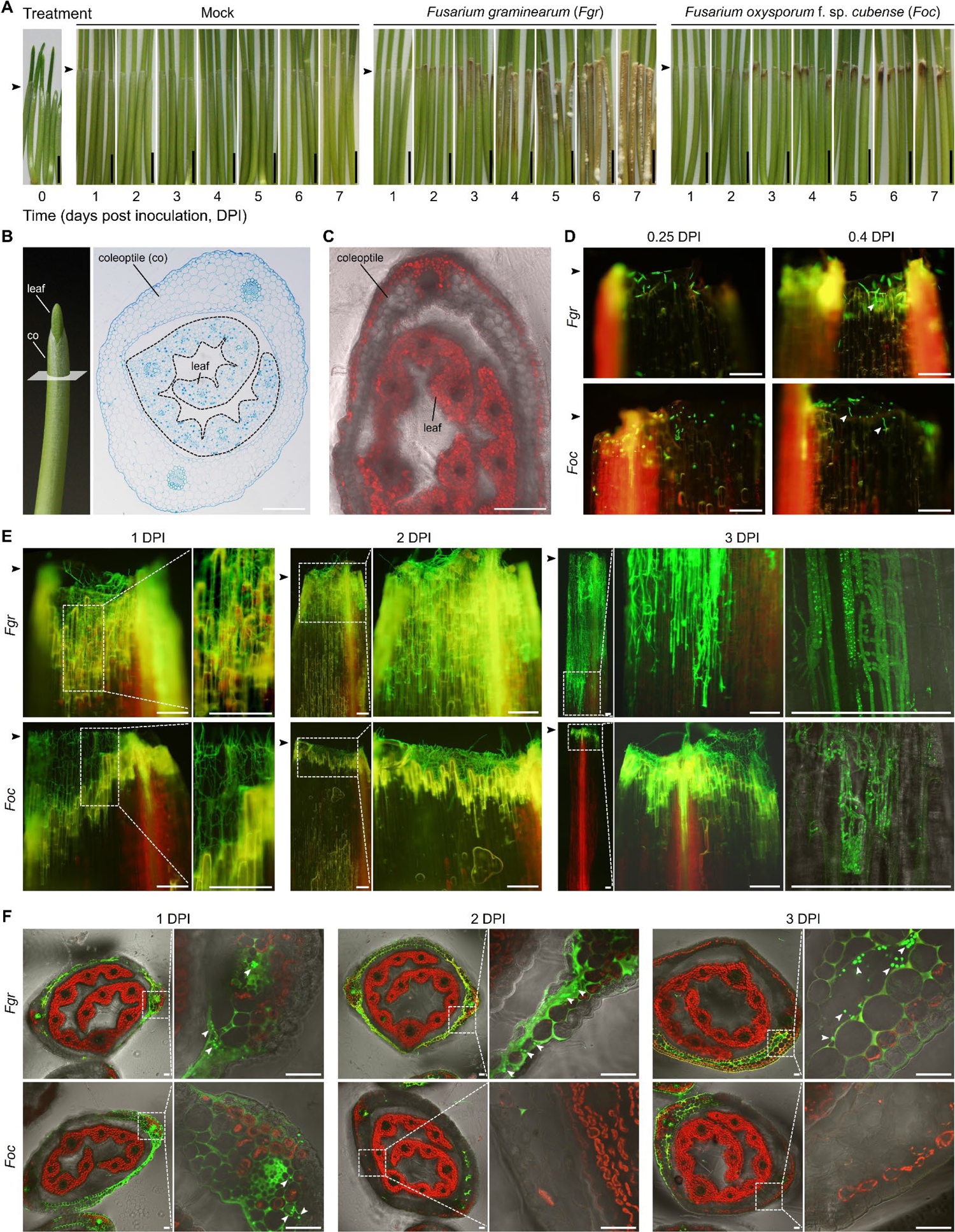
Invasion patterns of two *Fusarium* pathogens in wheat coleoptiles are similar in early stages, but diverge in later stages (A) Lesion size comparison in wheat coleoptiles ranging from day 0 to day 7 post-inoculation (DPI). The control group (Mock) involved inoculation with sterile water. *Fgr* group inoculated with *F. graminearum*. *Foc* group inoculated with *F. oxysporum* f. sp. *cubense*. A vertical black scale bar (5 mm) is present. Black arrowheads indicate the inoculation sites. (B) Structural overview of wheat shoots, complemented by a cross-section micrograph revealing the anatomical layout within the coleoptile. Young leaves, outlined by dashed lines, are shown enclosed by the coleoptile. Scale bar = 200 μm. (C) Chloroplasts, identifiable by their red auto-fluorescence, were found in mesophyll cells adjacent to the outer epidermis and those closest to the vascular bundles in the coleoptile. (D) (E) Periclinal microscopic observation of wheat coleoptiles inoculated by *Fgr* or *Foc* at indicated time points. The wheat coleoptiles exhibited auto-fluorescence of chloroplasts (red) in mesophyll cells adjacent to vascular bundles and of the plant cell wall (ranging from pale green to pale yellow). The position of the inoculation site in the coleoptile is labeled by black arrowheads. Both *Fgr* and *Foc* are transgenic strains, which can be identified by bright green fluorescence. The presence of germinated conidia is indicated by white arrowheads in (D). The advancing front of hyphae is shown in (E). Scale bar = 200 μm. (F) Transverse sections of coleoptiles near the advancing hyphal front, highlighting the distribution preferences of hyphae (bright green), as indicated by white arrowheads. The auto-fluorescence from chloroplasts (red) and plant cell walls (ranging from pale green to pale yellow) is also evident. Scale bar = 50 μm.

To compare the invasion progression of both *Fusarium* species in the wheat coleoptile, we used strains constitutively expressing GFP and conducted detailed cytological observations after inoculation. *In vitro*, the conidia of *Fgr* and *Foc* germinated after approximately 3 hours (Supplementary Figure 1A). *In planta*, within 1 DPI, conidia of both *Fusarium* species germinated at the wound sites on the coleoptiles (Figure 1D). Subsequently, extensive hyphal growth was observed within the layer of ruptured host cells around 1 DPI (Figure 1E). At 2 DPI, although *Fgr* penetrated the first one or two living cell layers, overall similar levels of hyphal growth were observed in both *Fusarium* species (Figure 1E). However, the divergence in invasion patterns between *Fgr* and *Foc* became pronounced at 3 DPI, where *Fgr* hyphae extended intra- or intercellularly far from the inoculation site, whereas *Foc* remained blocked at the wound of the coleoptile (Figure 1E).

As the host tissue of infection, the coleoptile possesses a symmetrical conical structure, with two vascular bundles on each flank, surrounded by mesophyll cells situated between the inner (adaxial) and outer (abaxial) epidermis (Figure 1B). One to three layers of mesophyll cells adjacent to the outer epidermis contain chloroplasts, while the remaining mesophyll cells are non-green (Figure 1C). In wheat seedlings of 3 to 6 days old (0 to 3 DPI), the coleoptile is in the cell elongation phase, during which the type and quantity of cells are largely fixed (Liptay and Davidson, 1972; Wiedenroth et al., 1990; Gibeaut et al., 2005). Microscopic observation of coleoptile serial cross-sections showed that the fluorescence of *Foc* and *Fgr* hyphae appeared at a similar distance from the line of inoculation sites before 2 DPI (Supplementary Figure 1B). Further microscopic observation revealed that *Fgr* hyphae rarely invaded the vascular bundles at the infection front (Figure 1F). Instead, *Fgr* preferentially invaded the non-green mesophyll with ample intercellular space, as evidenced in micrographs at 3 DPI (Figure 1F). Conversely, *Foc* exhibited uniform invading frequency in the vascular tissues and mesophyll cells of wheat coleoptiles (Figure 1F). Collectively, the invasion patterns of *Fgr* and *Foc* are similar in the early pathological stage up to approximately 2 DPI; subsequently, *Fgr* invasion speeds up, leading to the establishment of a susceptible interaction and successful colonization of the wheat coleoptile. In contrast, *Foc* invasion appears to halt and establish a non-host resistance.

### Establishment of scRNA-seq on wheat coleoptile infected by fungal pathogens from protoplast isolation to quality assessment

To investigate the single-cell transcriptional responses of the wheat coleoptile to *Fgr*, we designed an experiment incorporating a mock-treated group inoculated with sterile water as a negative control to avoid ambiguous stress responses (Figure 1A and 2A). Moreover, responses to the non-adapted pathogen *Foc*, used as a positive control in this setup, represent the effective nonhost resistance in the coleoptile (Figure 1A and 2A). Coleoptiles were collected just before treatment (i.e. 0 DPI) and at 1 to 3 DPI intervals (Figure 2A), corresponding to the primary colonization of *Fgr* on the coleoptile (Figure 1A). Viable protoplasts were efficiently isolated using the BD Rhapsody single-cell capturing platform (Figure 2B). High-quality data were obtained through Illumina sequencing, with sufficient depth to detect a robust gene expression profile per cell. On average, each library contained 4,623 cells and detected 82,387 genes, with an average of 4,082 genes detected per cell. Together, we employed a special case-control design and single-cell expression profiling pipeline to unveil wheat’s susceptibility to *Fgr*.

**Figure 2.**
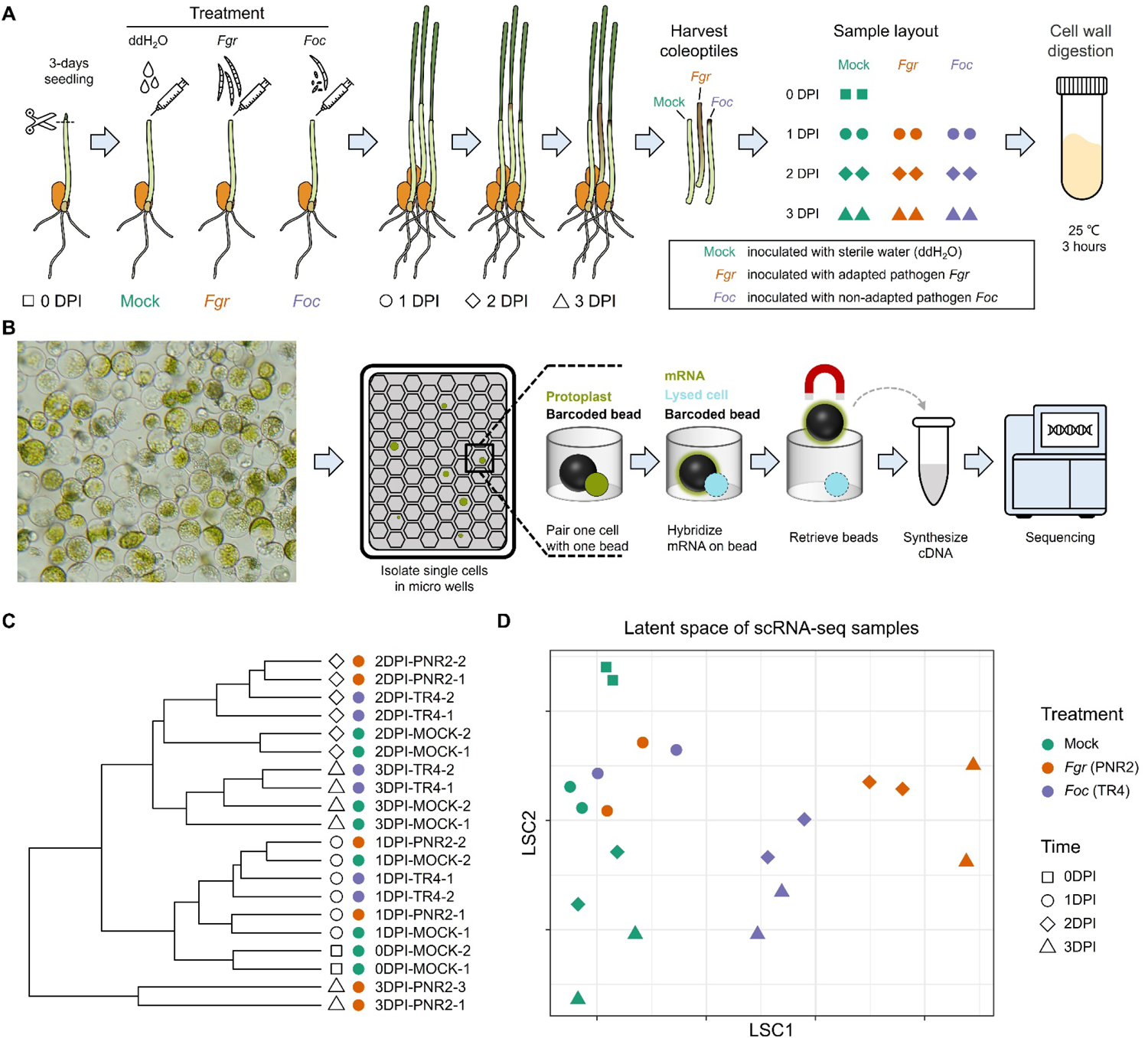
Establishment of scRNA-seq on wheat coleoptile from protoplast isolation to quality assessment (A) The schematic workflow depicts the experimental layout, encompassing sample collection and protoplast generation. Notably, samples collected before treatment are designated as ‘0DPI-MOCK’ to facilitate the analysis of sequencing data. Each sample type includes two biological replicates. (B) An overview of scRNA-seq library construction. The flowchart employs a green-filled circle to represent the coleoptile protoplast, while a black sphere symbolizes magnetic beads capturing mRNA (illustrated with a green gradient) from lysed cells (depicted as cyan-filled circles). (C) Hierarchical clustering of samples, based on Pearson’s correlation in averaged expression space, suggests the high reproducibility of the entire experiment. Samples collected from 0 to 3 DPI are represented by squares, circles, diamonds, and triangles, respectively. Various treatments are indicated by different colors. The cladogram’s leaves are labeled with actual sample names. PNR2 represents the *Fgr*-treatment, and TR4 represents the *Foc*-treatment. (D) Visualization of multiple single-cell samples in latent space reveals that quality control and batch effect removal procedures effectively preserve biological variance. LSC1/2 refers to latent space components 1 and 2, respectively. This latent space is calculated based on the Wasserstein Distance between ‘cell clouds’ in the PCA space of single-cell expression profiles.

Single-cell RNA sequencing of multiple samples can include low-quality cells and batch effects, which may hinder downstream analysis. Adhering to the three criteria outlined in the Methods section, we excluded 1147 (1.21%) cells of low quality, followed by the removal of ambient RNA contamination and batch effects. Consequently, 91,320 cells were retained for further analysis. To evaluate the reproducibility of biological replicates, we performed hierarchical clustering, visualized in a cladogram (Figure 2C). This analysis revealed that samples from the same time point and treatment were grouped together, with minimal distance between replicates (Figure 2C). To assess the preservation of biological variance post-quality control and batch effect removal, we projected the samples onto a latent space, with distances reflecting differences (Figure 2D). The scatter plot demonstrates that sample variations were mainly attributable to biological factors including time and treatment (Figure 2D). In summary, the applied quality control and batch effect removal procedures effectively preserved biological variance within the samples.

### Identification and characterization of wheat coleoptile cell types by combined single-cell and spatial transcriptomics

To avoid interference of biotic stress on cell type identification, we first identify cell types in mock-treated samples and then predict their identities in pathogen-challenged samples to obtain an integrated dataset. Initially, cells were grouped via unsupervised clustering based on their expression profiles. The number of clusters is influenced by resolution, where higher values result in a greater number of clusters. Optimal clustering was achieved at a resolution of 0.6, which prevented over-clustering and yielded cell clusters whose number was consistent with the anticipated complexity of the wheat coleoptile (Supplementary Figure 2A). Consequently, cells from mock samples, ranging from 0 to 3 DPI, were categorized into 15 clusters (Figure 3A).

**Figure 3.**
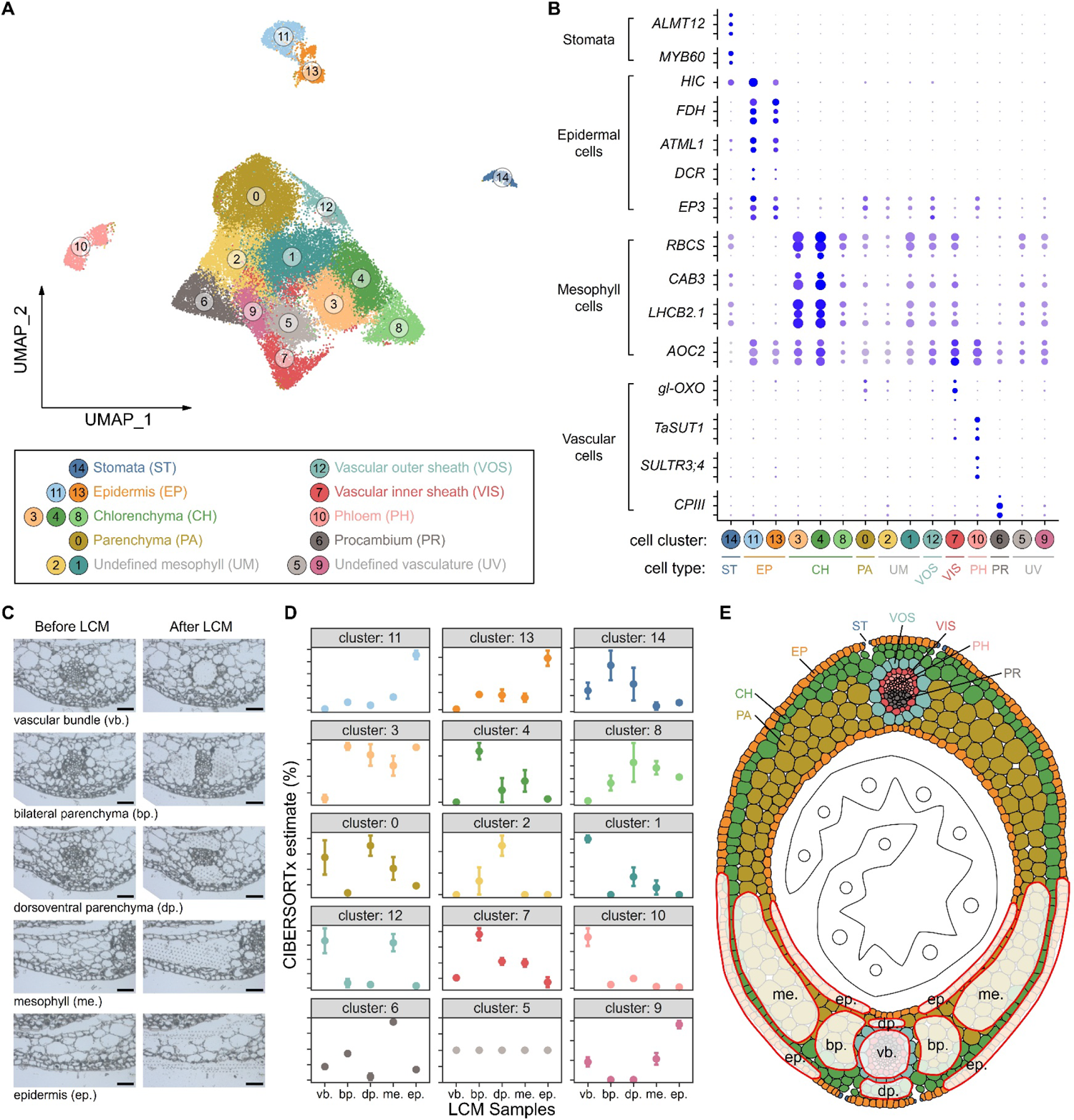
Identifying wheat coleoptile cell types using combined single-cell and spatial transcriptomics (A) UMAP projection of cells from Mock-treatment samples, where each cell is represented as a dot and color-coded by cluster. Cells are categorized into 15 distinct clusters based on their expression profiles, with each cluster being assigned a specific cell type, as indicated in the black box in the lower panel. (B) Expression patterns of known marker genes or their wheat homologs. Triplets of homoeologues from wheat sub-genomes are grouped together on the y-axis and labeled concisely. The dot size represents the proportion of cluster cells expressing a particular gene, while the color denotes the average gene expression level in a cell cluster. Detailed information on selected genes, including IDs and references, is provided in Supplementary Table 1. (C) Illustration of tissues selected for laser capture microdissection (LCM), identifiable by blank areas with charred spot arrays, evident when comparing micrographs before and after LCM. The scale bar represents 50 μm. (D) Estimated abundance of different cell clusters in anatomically distinct LCM samples. Displayed are the mean values derived from CIBERSORTx estimates, accompanied by error bars indicating the standard error of the mean. Each LCM sample was biologically replicated thrice (n = 3). Abbreviations for sample names are detailed in panel (C). (E) A schematic representation of the anatomy and cell types of a wheat coleoptile in cross-section. The lower half of this diagram highlights selected areas for LCM (translucent white polygons with red borders). Abbreviations for cell types and LCM samples are referenced in panels (A) and (C).

Wheat seedling coleoptile comprises cells derived from three body layers: layer 1 (L1), constituting a single-cell layer of epidermis; layer 2 (L2), which is the mesophyll; and layer 3 (L3), the site of vascular initiation (Zeng et al., 2016). To assign identities to the 15 cell clusters, we first supplemented spatial information obtained from RNA sequencing of laser captured cell samples (LCM-based RNA-seq). According to the anatomy of wheat coleoptiles, the vascular bundle (L3), the bilateral and dorsoventral parenchyma cells adjacent to the vascular bundle (L2), mesophyll cells (L2), and epidermis (L1) (Figure 3C and the lower half of Figure 3E) were isolated by LCM and subjected to bulk RNA sequencing. Expression profiles of LCM samples were decomposed based on single-cell transcriptomic data of the wheat coleoptile using a deconvolution method, CIBERSORTx (Newman et al., 2019). The proportions of cell clusters in each LCM sample revealed their spatial allocation (Figure 3D), thereby directly elucidating the body layers in which these cell clusters are allocated, including the L1 epidermal cell clusters 11 and 13, the L2 various mesophyll cell clusters 3, 4, 8, and 2, and L3 vascular clusters 12 and 10.

After most cell clusters were spatially assigned to the three body layers, we further used a set of well-annotated marker genes with known expression patterns as indicative of cell identity (Supplementary Table 1). This resulted in the identification of the phloem (cluster 10), procambium (cluster 6), vascular inner sheath (cluster 7), epidermis (clusters 11 and 13) and stomata (cluster 14). The mesophyll, constituting the largest proportion of the coleoptile, exhibits variation in morphology and function, but lacks a clear separation of different cell subpopulations. Therefore, the continuum of mesophyll cells in the L2 layer was categorized *in silico* into three cell types based on gene expression or RNA *in situ* hybridization assays: chlorenchyma (clusters 3, 4, 8), parenchyma (cluster 0), and vascular outer sheath (cluster 12). Further details on the identification of wheat cell types are provided in Supplementary Note 1.

Through cell abundance and gene expression analysis detailed in Supplementary Note 2, we observed that developmental features of the eight cell types elucidate the maturation and senescence of wheat coleoptile under the mock treatment. In the final step of the cell type identification approach (see Methods), we used the annotated mock dataset as a reference for prediction and integrated the transcriptomes of cells from Mock, *Fgr*, and *Foc* treatments into a unified transcriptional space, collectively identifying eight cell types (Figure 4A and 4B).

**Figure 4.**
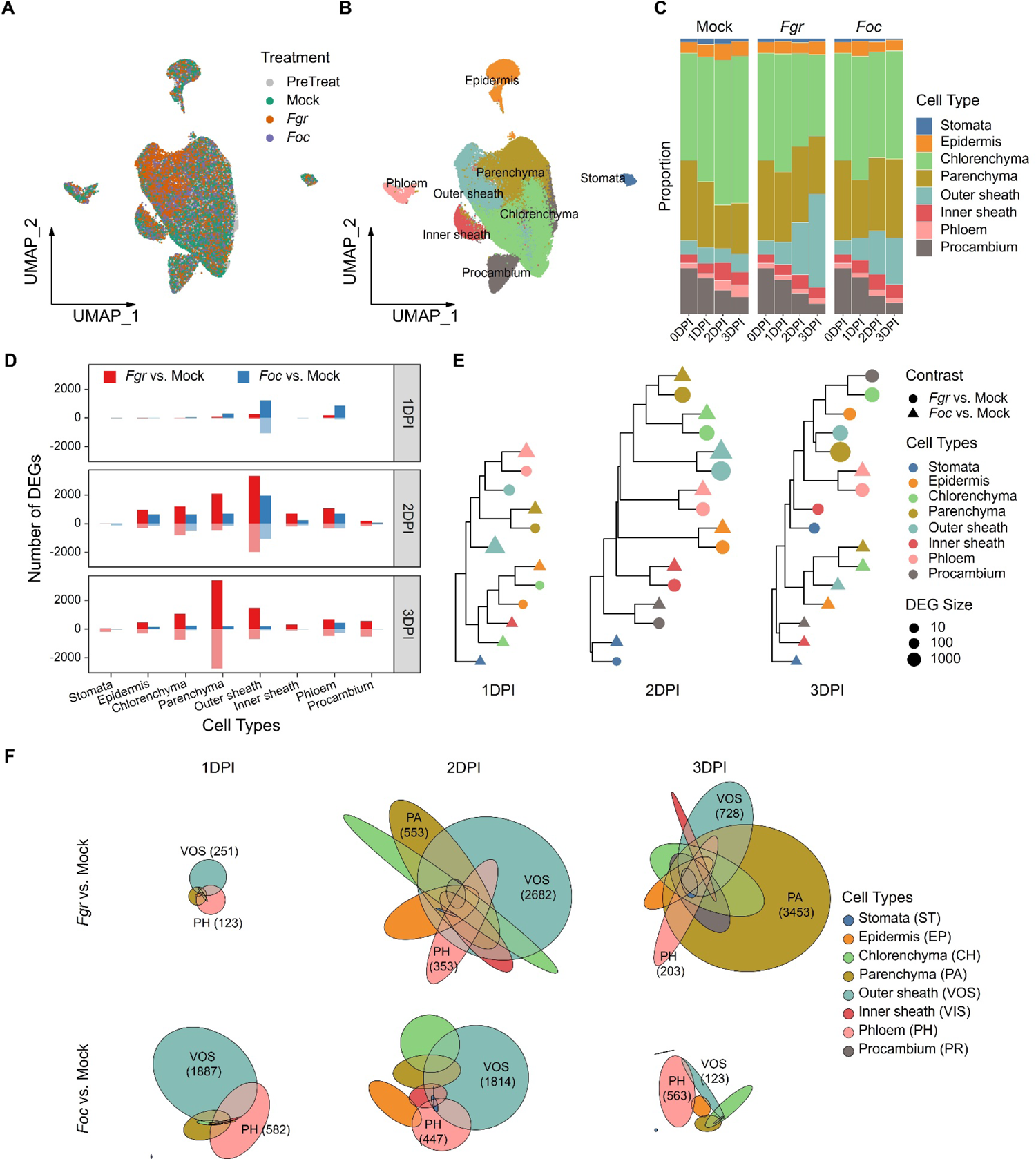
The cellular heterogeneity of wheat coleoptiles in response to *Fusarium* infection was observed both quantitatively and qualitatively (A) The transcriptomes of cells from Mock, *Fgr*, and *Foc* treatments were integrated into a unified transcriptional space. Cells in the UMAP projection are depicted as dots, color-coded by treatment. Mock represents cells inoculated with sterile water. *Fgr* denotes cells inoculated with *F. graminearum*, and *Foc* with *F. oxysporum* f. sp. *cubense*. PreTreat refers to cells from pre-treatment samples, also known as ‘0DPI-MOCK’. (B) Visualization of eight cell types, analogous to (A), with dots color-coded by cell types. These types were predicted and transferred from annotated Mock-treatment datasets. (C) Average proportion of cell types within each sample group. (D) The quantification of differentially expressed genes (DEGs) highlights the varying capacities of cell types to respond to *Fgr* and *Foc*. Semi-transparent bars represent down-regulated genes, indicated by negative values. (E) Hierarchical clustering of DEG sets to elucidate the similarities and differences in cellular responses to pathogens across cell types. The distance between sets was calculated using Jaccard’s index. The cladogram leaves are represented by points, encoding DEG sets with shapes indicating contrast, colors signifying cell types, and sizes correlating with the number of DEGs. This hierarchical clustering was conducted separately for 1 to 3 DPI. (F) Area-proportional Euler diagrams of DEG sets for each treatment group, visually depicting the overlap and distinction in cell type responses. Numbers in brackets specify the count of DEGs in each subset.

### Cell types of the coleoptile differed in their capacity to respond to pathogen infection, which was constrained by cell identity

Next, we sought to resolve the transcriptional reprogramming of the coleoptile and investigate the capacity of different cell types to respond to pathogen infection. Using the mock treatment as a control, we conducted differential expression analyses for each cell type in response to *F. graminearum* (*Fgr* vs. Mock) or *F. oxysporum* f. sp. *cubense* (*Foc* vs. Mock) infection, based on pseudo-bulk expression profiles. Thereby, we established a transcriptome atlas of the wheat coleoptile in response to *Fgr* and *Foc* (Figure 4B and Supplementary Table 2). Evaluating the number of differentially expressed genes (DEGs) as an indicator of the amplitude of stress response, we observed that the stomata and procambium showed minimal responses to fungal invasion (Figure 4D). Conversely, the outer sheath and phloem exhibited the most pronounced responses to fungal invasion across 1 to 3 DPI (Figure 4D).

To delineate how the host cell’s response is influenced by its identity, we utilized DEG sets as proxies for cell-type-specific responses. Our analysis revealed that the outer sheath, phloem, and parenchyma displayed the most distinct responses to *Fusarium* invasion in the coleoptile (Figure 4F). We defined the similarity between DEG sets based on the size of the intersection relative to the union of these sets.

In the cladogram resulting from hierarchical clustering, which groups similar responses, DEG sets were predominantly clustered according to their cell types rather than treatments during 1 to 2 DPI, except for those with very low sizes at 1 DPI (Figure 4E). However, with the exception of the phloem, responses of cell types at 3 DPI were primarily grouped by treatments (Figure 4E). This pattern indicates that wheat’s responses to pathogens, initially constrained by cell identity during 1 to 2 DPI, were eventually overridden by susceptible interactions with *Fgr* at 3 DPI, altering the developmental features of each cell type except for phloem.

### Responsiveness of the phloem and outer sheath in the rapid onset of nonhost resistance was suppressed in early stage of the susceptible interaction

Subsequently, we further investigated the differences between wheat’s nonhost resistance to *Foc* and its susceptible interaction with *Fgr*. The latent space plot presents sample-level transcriptomic similarity, showing that coleoptiles under various treatments were not apparently different at 1 DPI (Figure 2D). By 2 DPI, the coleoptiles treated with two *Fusarium* species were similar but differed from the mock treatment (Figure 2D). However, at 3 DPI, the coleoptiles treated with the non-adapted pathogen *Foc* were more similar to the mock treatment than those treated with the adapted pathogen *Fgr* (Figure 2C and 2D). The numbers of DEGs further illustrate the transcriptional reprogramming of coleoptiles under different treatments. Wheat’s nonhost resistance to *Foc* commenced at 1 DPI, peaked at 2 DPI, but rapidly diminished at 3 DPI (Figure 4D). In contrast, the number of DEGs induced by the adapted pathogen *Fgr* was very low but increased drastically at 2 DPI and remained high through 3 DPI (Figure 4D). Taken together, the nonhost resistance to *Foc* was mounted rapidly and limited in amplitude, while the responses to *Fgr* were delayed during 1 to 2 DPI in comparison to those to *Foc* and burst at 3 DPI.

In wheat’s nonhost resistance to *Foc*, the outer sheath and phloem are the two most rapidly and drastically responsive cell types. At 1 DPI, the expression of 1219 and 584 genes, respectively in these cell types, was significantly increased in comparison to mock samples (Figure 4D and 4F). Genes in chitin catabolism and hydrogen peroxide catabolism are significantly enriched in the up-regulated DEGs of all three cell types, suggesting that degradation of fungal cell wall and generation of reactive oxidative species are common in the first wave defense of these three cell types (Supplementary Figure 5B). The outer sheath’s specific responses at 1 DPI included upregulated expression of genes in chorismate biosynthesis, glutathione metabolism, and arginine biosynthesis, along with suppressed expression of genes in photosynthesis (including photosystem II stabilization, electron transport, and photosynthesis dark reaction), response to heat, and water transport (Supplementary Figure 5B). This suggests that biosynthesis of defense hormone salicylic acid, increase of buffer capacity of oxidative burst, reduction of photosynthesis, and reduction of water fluidity might be rapid defense approaches allocated to the outer sheath. In contrast, the phloem exhibited a distinct increase in the expression of genes involved in defense responses to fungi, cell wall biogenesis, and negative regulation of proteolysis under *Foc* treatment at 1 DPI (Supplementary Figure 5B), suggesting a distinct first wave defense strategy in the phloem including cell wall reinforcement and protection from proteases (probably from fungus). Although a greater number of genes were induced or suppressed in the outer sheath during 1 to 2 DPI, quantitatively, the phloem’s nonhost resistance proved more sustainable than that of the outer sheath, as evidenced by the drastic decrease in DEGs in the outer sheath at 3 DPI (Figure 4D and 4F). Excluding stomata and procambium, which showed minimal response to the non-adapted pathogen *Foc*, other cell types exhibited moderate nonhost resistance, predominantly at 2 DPI (Figure 4D and 4F). Collectively, these results underscore the overlapping yet divergent behaviors of the outer sheath and phloem in the rapid onset of nonhost resistance to *Foc*.

In the susceptible interaction between wheat and *Fgr*, the invasion progress did not differ from that of *Foc* at 1 DPI (Figure 1E and 1F). However, the outer sheath and phloem showed diminished response amplitudes to *Fgr*, especially compared to that triggered by the non-adapted pathogen *Foc* (Figure 4D). It’s worth noting that the phloem’s response to both *Fgr* and *Foc* was strikingly similar during 2 to 3 DPI, as evidenced by the comparative number and overlap of DEG sets (Figure 4D, 4E, and Supplementary Figure 5A). This similarity was further supported by enrichment analyses of DEG sets induced or suppressed by *Fgr* and *Foc* in the phloem (Supplementary Figure 5C and 5D). Interestingly, genes involved in the glutathione metabolic process, specifically encoding glutathione S-transferases, were induced by *Fgr*, yet suppressed by *Foc* from 1 to 3 DPI (Supplementary Figure 5B, 5C, 5D, and Supplementary Table 3). The similar basal defense of the phloem during 2 to 3 DPI highlights the significance of suppressing the phloem’s responsiveness in the susceptible interaction with *Fgr* at 1 DPI.

For the outer sheath, responses to the adapted pathogen *Fgr* during 2 to 3 DPI largely overlapped with nonhost resistance during 1 to 2 DPI, but also included a substantial number of unique DEGs (Supplementary Figure 5A). Noteworthy, among the 1219 DEGs that were upregulated significantly at 1 DPI under non-adapted *Foc* infection in the outer sheath, only 194 (16%) were also upregulated significantly at 1 DPI upon *Fgr* infection. However, 655 (54% of 1219) DEGs that were not upregulated at 1 DPI exhibited a significant increase in expression at 2 DPI or 3 DPI in the outer sheath upon *Fgr* infection, indicating a delayed response to the adapted pathogen *Fgr* compared to the non-adapted pathogen (Supplementary Table 2). Enrichment analysis revealed that the induction of biological processes by *Fgr* in the outer sheath was delayed until 2 DPI, in contrast to the rapid onset of nonhost resistance. These delayed processes include biosynthesis of chorismate, arginine, tryptophan, and polyamines; L-phenylalanine catabolism; S-adenosylmethionine metabolism; and glutathione metabolism (Supplementary Figure 5B, 5C, 5D, and Supplementary Table 3). Similarly, the suppression of various biological processes by *Fgr*, including photosynthesis, regulation of protein stability, water transport, hydrogen peroxide catabolism, and responses to heat and hydrogen peroxide, was also delayed to 2 DPI (Supplementary Figure 5B, 5C, 5D, and Supplementary Table 3). Additionally, various cell types within the coleoptile shared a lot of similar biological processes that were differentially regulated in response to *Fgr* infection at 3 DPI (Supplementary Figure 5D). In summary, the responsiveness of the outer sheath was also suppressed in the susceptible interaction with *Fgr* at 1 DPI, in contrast to nonhost resistance.

### Transcriptional activation of immune pathways was compartmentalized in nonhost resistance, but widespread in susceptible interaction

Due to the limitations and ambiguities in the Gene Ontology (GO) annotation of the wheat genome, we manually curated a list of genes associated with defense pathways (Supplementary Table 4), grounded on the current understanding of the plant immune system (Ngou et al., 2022). As depicted in Figure 5A heatmaps, each row illustrates the expression level of a single gene across eight cell types, with treatments—mock-inoculated (green), adapted *Fgr*-inoculated (red), and non-adapted *Foc*-inoculated (blue)—displayed from left to right within each cell type column. The heatmaps also list time points (0-, 1-, 2-, and 3-DPI) within each treatment block.

**Figure 5.**
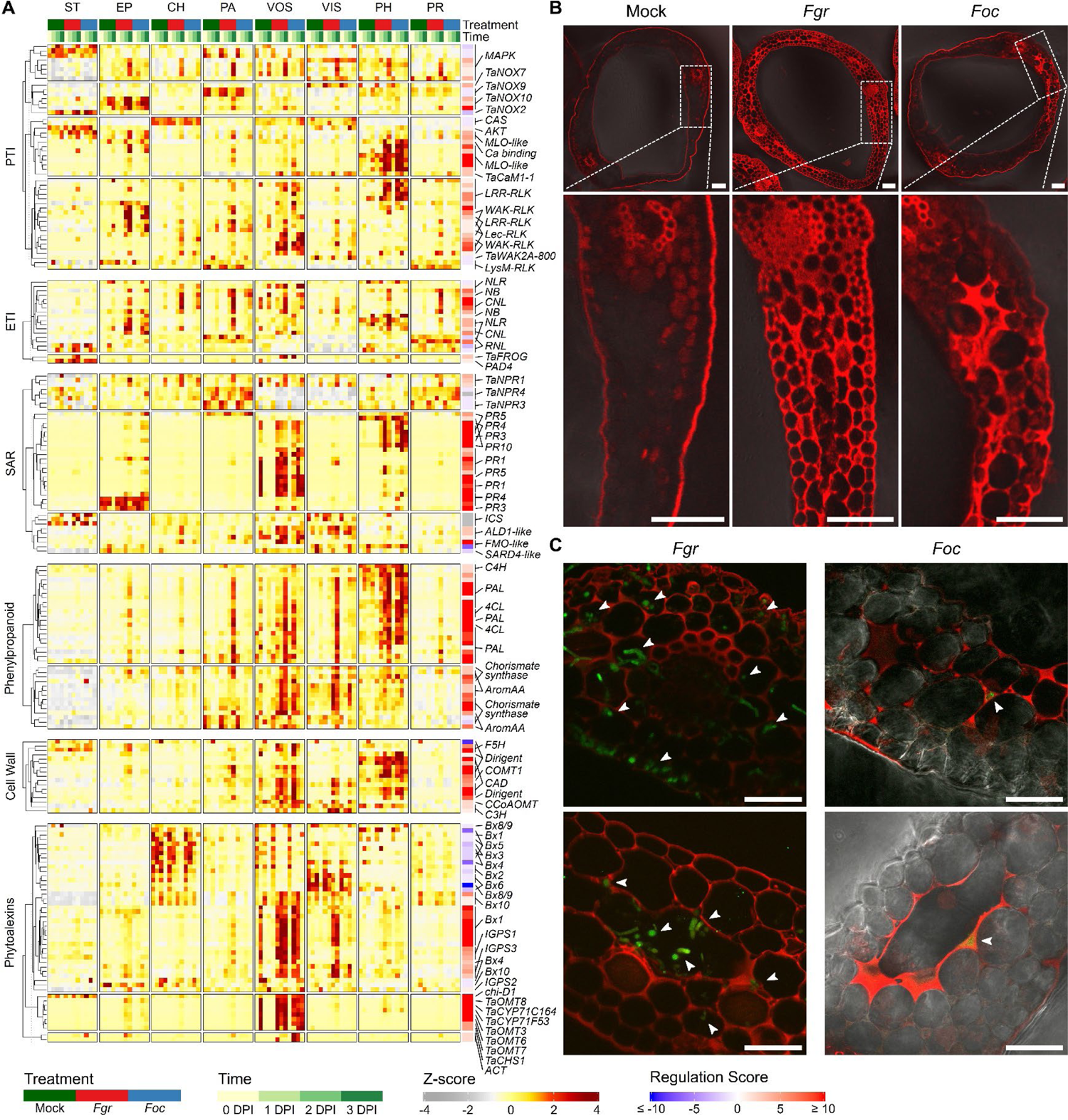
The cell-type-specific upregulation of defense genes reveals the compartmentalization of immunity and secondary metabolism in the wheat coleoptile (A) The heatmap displays the expression level (Z-score) of defense genes across eight cell types under *Fusarium* infection. The columns of the heatmap were split from left to right into Stomata (ST), Epidermis (EP), Chlorenchyma (CH), Parenchyma (PA), Outer sheath (VOS), Inner sheath (VIS), Phloem (PH), and Procambium (PR). Treatments are shown from left to right within each cell type column, including mock-inoculated (green), adapted *Fgr*-inoculated (red), and non-adapted *Foc*-inoculated (blue). The heatmaps also show four distinct time points (0-, 1-, 2-, and 3-DPI) within each treatment block. The Regulation Score is displayed in a separate column to the left of the row names, using blue and red to indicate down- and up-regulation of genes in the coleoptiles. (B) After the non-adapted pathogen *Foc* infection at 3 DPI, the parenchyma exhibited dense, mosaic-like lignin deposition in localized areas. Scale bars = 100 μm. (C) Lignin deposition (red fluorescence) effectively blocked the hyphae of *Foc* but failed to prevent the adapted pathogen *Fgr* from spreading. The presence of fungal hyphae (green fluorescence) is indicated by white arrowheads. Scale bars = 50 μm.

A detailed expression pattern analysis of PTI, ETI, and SAR-related genes (Figure 5A and Supplementary Note 3) revealed that non-overlapping PRRs and PTI signaling components including calcium channels, MAPKs, ROS producers, and others, were variously induced in the epidermis, outer sheath and phloem. This finding indicates that coleoptile cells vary in their pathogen-sensing abilities. The epidermis and phloem cells exhibited a roughly complete PTI pathway compared to the outer sheath. However, in contrast to the rapid and cell-type-specific nonhost resistance of wheat to *Foc*, the upregulation of PTI recognition (i.e. PRRs) and signaling components occurred at mismatched times and in inadequate cell types during the susceptible interaction with *Fgr*. Similarly, the limited upregulation of ETI (i.e. NLRs) in nonhost resistance was boosted to more cell types during the susceptible interaction with *Fgr* at 3 DPI.

Regarding SAR, the upregulation of signaling molecule biosynthesis pathways (salicylic acid and N-hydroxypipecolic acid) in nonhost resistance to *Foc* was mainly confined to the phloem and outer sheath, but adapted pathogen *Fgr* induced a stronger and more widespread upregulation at a later stage (Figure 5A and Supplementary Note 3). Additionally, *Fgr* infection significantly enhanced the perception of SAR signaling in more cell types during 2 to 3 DPI, through the upregulation of activators (*NPR1*) and downregulation of repressors (*NPR3*/*NPR4*). The upregulation of pathogenesis-related (PR) proteins-encoding genes (PR-1, chitinase, thaumatin-like proteins, and Bet v1-like proteins) was restricted to the phloem and outer sheath, with no marked differences between adapted and non-adapted *Fusarium* infections (Figure 5A and Supplementary Note 3).

Secondary metabolites play a crucial role in plant immune responses by acting as signaling molecules and defensive compounds against pathogens and herbivores. Therefore, we analyzed the expression patterns of biosynthetic genes for several phytoalexins, including isoflavones, hydroxycinnamic acid amides (HCAAs), and benzoxazinoids (BXDs) during fungal infections (Figure 5A and Supplementary Note 4). The upregulation of isoflavone biosynthesis was partitioned within two spatially isolated cell types, with precursors in the phloem and final products in the outer sheath. Under mock treatments, the biosynthesis and storage of BXDs were organized in two distinct cell types. However, *Fusarium* infection enhanced the production of BXD precursors in a third cell type, the outer sheath (Figure 5A and Supplementary Note 4). The biosynthesis of HCAAs was specifically induced in the outer sheath following infection of the non-adapted pathogen *Foc*, whereas *Fgr* prevented the upregulation of relevant biosynthetic genes to reduce the production of HCAA (Figure 5A and Supplementary Note 4).

Downstream responses of PTI in plants can lead to cell wall reinforcement (Miedes et al., 2014). Lignin, a heterogeneous polymer of monolignols, strengthens the cell wall against pathogens. In wheat’s nonhost resistance to *Foc*, the transcriptional activation of lignin monomer biosynthesis from phenylalanine was more integrated in the outer sheath than in the phloem (Figure 5A and Supplementary Note 4). However, the infection of *Fgr* impaired the polymerization of lignin monomers by diminishing or delaying the expression of dirigent proteins in the outer sheath (Figure 5A and Supplementary Note 4).

To validate lignin deposition in coleoptiles infected by *Fusarium* pathogens, Basic Fuchsin, a fluorescent dye for lignin, was employed to stain serial hand cross-sections. In mock samples from 1 to 3 DPI, lignin deposition was exclusively observed in the phloem, inner sheath, and epidermis (Figure 5B). Following the non-adapted pathogen *Foc* infection, dense, mosaic-like lignin deposition appeared locally in the parenchyma (Figure 5B). In contrast, upon adapted pathogen *Fgr* infection, loose lignin deposition filled nearly all intercellular spaces by 3 DPI (Figure 5B). To confirm that lignin deposition effectively prevented pathogen spread, the fluorescence of Basic Fuchsin and the hyphae of *Fusarium* carrying green fluorescent proteins were simultaneously examined. In the parenchyma, lignin was locally deposited around the hyphae of the non-adapted pathogen *Foc*, with no hyphae detected beyond this region (Figure 5C). However, despite widespread lignin deposition in the parenchyma, the hyphae of the adapted pathogen *Fgr* extensively spread (Figure 5C). In summary, loose and widespread lignin deposition did not prevent the spread of the adapted pathogen *Fgr*, in contrast to the dense and local lignin deposition induced by the non-adapted pathogen *Foc*. These observations suggest that *Fgr* may breach the cell wall reinforcement by delaying lignin polymerization.

### Parenchyma cells underwent a transition to divergent immune-activated states after exposure to different fungal pathogens

The wheat-*Fusarium* interaction within the coleoptile represents a complex spatiotemporal dynamic, particularly at the advancing infection front where only certain host cells directly contact *Fgr* hyphae (Zhang et al., 2012). Cytological observations revealed that *Fgr* hyphae preferentially proliferated in the intercellular spaces of parenchyma at the infection front (Figure 1F). It is worth noting that the proportion of parenchyma and outer sheath cells underwent significant changes over time in pathogen-infected samples, with a decrease in the parenchyma cells and an increase in the outer sheath cells (FDR < 0.05) (Figure 4C and Supplementary Figure 4E). In line with this, when projecting coefficients from co-varying neighborhood analysis (Reshef et al., 2022) on UMAP embeddings, results suggest that the expansion of chlorenchyma in the coleoptile of mock samples was substituted by an expansion of outer sheath over time under fungal treatments (Supplementary Figure 7D). In other words, in the transcriptome continuum of mesophyll, the proportion of parenchyma and outer sheath cells underwent a significant decrease and increase, respectively, in response to fungal infection.

During the data integration of multiple coleoptile samples, two subpopulations of the outer sheath, clusters 7 and 12, are notable for their low prediction confidence, differing from conserved cell types like stomata, epidermis, and phloem (Supplementary Figure 7C and 4D). This suggests a classification of many cells exposed to pathogen infection as outer sheath, based solely on transcriptomic similarity under biotic stress, irrespective of their actual cell identity. RNA velocity analysis calculates the relative abundance of unspliced and spliced mRNA from sequencing data (La Manno et al., 2018), which can be used in predicting the future transcriptional states of individual cells on a timescale of hours. RNA velocity vectors of L2-derived cells were projected on UMAP embeddings (Supplementary Figure 7B). Results show that the future transcriptional states of parenchyma cells are similar to the outer sheath in response to pathogen treatment. Furthermore, the outer sheath exhibited the highest capacity in response to pathogens, enriching immunity-related genes (Figure 4D and 5A). Hence, these findings from RNA velocity and cell abundance analysis suggest that parenchyma cells may acquire transcriptional states of the outer sheath after exposure to fungal invasion.

To explore the temporal dynamics of these pathogen-responsive mesophylls, we applied pseudotime analysis to the parenchyma and outer sheath cells. A tree-like trajectory was fitted in the transcriptional space of these cells, and pseudotime values which indicate the defense progression, were assigned to each cell in the trajectory (Figure 6A and Supplementary Figure 7E). Branches of the trajectory, which diverged from a common upstream in parenchyma cells, culminated in distinct terminal cell states within the outer sheath (Figure 6A and Supplementary Figure 7E). The distribution of inferred pseudotime aligned roughly with actual sampling time points (Supplementary Figure 7F). Differential abundance analysis between pathogen treatments and mock samples revealed distinct terminal states enriched with cells exposed to adapted and non-adapted *Fusarium* pathogens (Figure 6A and 6B). Thus, each trajectory branch represents a distinct defense strategy of the pathogen-responsive mesophyll, with branches 1 and 3 representing a common defense strategy against both adapted and non-adapted *Fusarium*, and branch 2 indicative of an *Fgr*-specific defense strategy.

**Figure 6.**
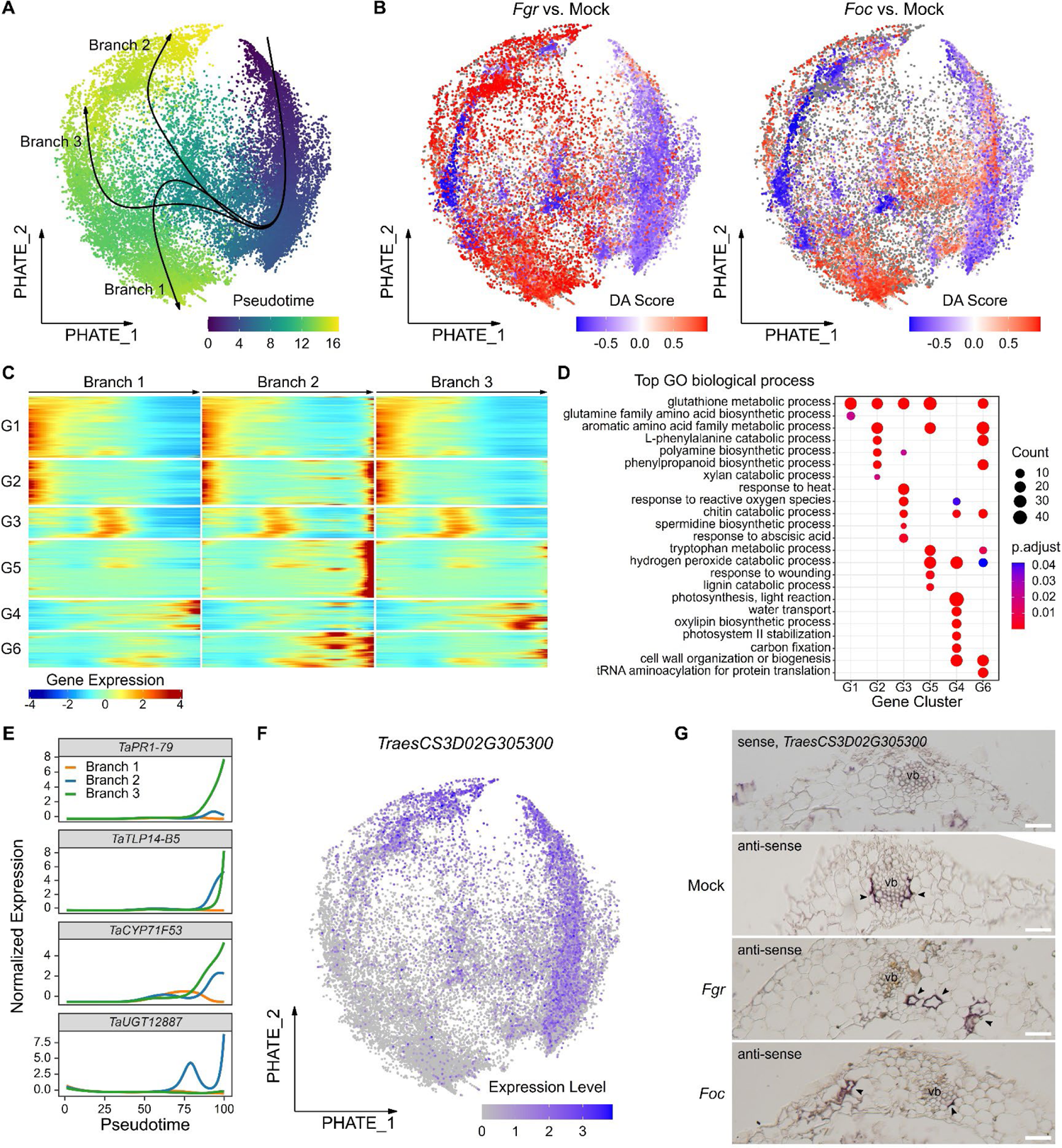
Gradual transition of parenchyma cells to divergent immune-activated states upon *Fusarium* infection (A) A tree-like trajectory illustrates the smooth transition of parenchyma cells to varied terminal states during continuous interactions with *Fusarium* pathogens. Parenchyma and outer sheath cells are visualized using PHATE projection, with fitted principal curves shown in black to represent these transitions. Each point, denoting a cell, is colored by its average pseudotime, indicating transcriptional progression towards a terminal state. (B) Differential abundance analysis reveals that cells exposed to adapted and non-adapted *Fusarium* infection were enriched in different trajectory branches. Differential abundance analysis was performed by contrasting fungi-invaded groups (*Fgr* or *Foc*) from 1 to 3 DPI against a control group (Mock) across 0 to 3 DPI. Each dot in the PHATE projection represents a cell, color-coded by the Differential Abundance Score (DA Score), where larger (red) values indicate higher abundance in *Fgr* or *Foc* samples, and smaller (blue) values signify Mock samples. (C) A heatmap shows the expression patterns of genes differentially expressed along the trajectory, with arrows marking the direction. These genes are categorized into six modules (G1 to G6) using k-means clustering based on their expression patterns. (D) Gene Ontology enrichment analysis of gene modules identified in (C) reveals the regulatory divergence in biological processes during interactions with adapted and non-adapted *Fusarium* pathogens. (E) The continuous interactions of *Fusarium* pathogens with parenchyma cells along different trajectory branches led to the upregulation of representative immune marker genes. (F) A secretory peroxidase-encoding gene differentially expressed along branch 2 was selected as a marker of the *Fgr*-enriched interaction. (G) RNA *in situ* hybridization confirms the expression of this marker gene in the parenchyma cells upon *Fusarium* infection. Red arrowheads mark the appearance of transcripts. vb, vascular bundle. Scale bar = 50 μm.

To elucidate the biological significance of parenchyma defense strategies, we applied a generalized additive model (Van den Berge et al., 2020) based on Mock, *Fgr*, and *Foc* treatments to investigate gene expression dynamics across different trajectory branches. Expression analysis of known defense genes along pseudotime (Supplementary Figure 8 and Supplementary Note 5) revealed a transcriptional upregulation of secretory peroxidases (*TaPOD237* and *TaPOD363*) in branches of the parenchyma trajectory. The peroxidase-mediated chemotropism of *Fgr* (Sridhar et al., 2020; Sridhar et al., 2023) indicates a spatial association between fungal hyphae and parenchyma cells. As general features of parenchyma defense strategies, pathogenesis-related genes (*PR1*s and *TaTLP*s), Fusarium head blight-resistance genes (*TaUGT12887* and *TaSAM*), and defensive secondary metabolite biosynthesis (e.g. *TaCYP71F53*) were upregulated along branches of the parenchyma trajectory, but with varying amplitude in each branch (Supplementary Figure 8 and Supplementary Note 5). However, certain genes (*TaACT*, *TaFROG*, and some uncharacterized genes) exhibited treatment-specific upregulation patterns, being exclusively upregulated under either *Foc* or *Fgr* treatment (Supplementary Figure 8 and Supplementary Note 5). This finding reveals the competition between immune and susceptible processes in the parenchyma cell state transitions.

To further characterize the difference between common and *Fgr*-specific defense strategies of the parenchyma cells, we identified genes associated with pseudotime in different trajectory branches and clustered them into six modules based on expression patterns (Figure 6C and Supplementary Table 5). Gene Ontology enrichment analysis of these gene modules revealed representative pathways of the parenchyma cell state transitions. Modules G1 and G2, present in all branches, exhibiting gradual downregulation along the pseudotime (Figure 6C), were enriched with the biosynthesis of glutamine, polyamine, and phenylpropanoid (Figure 6D). Module G3, showing transient upregulation in all branches, was enriched with response to heat, reactive oxygen species, and abscisic acid, as well as the chitin catabolism and spermidine biosynthesis, indicating basal immune responses to both *Fusarium* species (Figure 6D). Module G4, gradually upregulated in common defense strategy (branches 1 and 3), contrasted with modules G5 and G6, which indicated late and gradual upregulation in branch 2, characteristic of the *Fgr*-specific strategy (Figure 6C). Modules G4, G5, and G6 were enriched with pathways like hydrogen peroxide catabolism, cell wall organization, and chitin catabolism (Figure 6D). However, the common defense strategy was further defined by the gradual upregulation of photosynthesis, water transport, and oxylipin biosynthetic processes in module G4 (Figure 6D). Conversely, the metabolism of glutathione, phenylpropanoid, and tryptophan, along with the lignin catabolic process and response to wounding, were enriched in modules G5 and G6, characteristic of the *Fgr*-specific strategy (Figure 6D). In summary, these results suggest that the regulatory divergence of the immune-related pathways, particularly photosynthesis and phenylpropanoid metabolism, drives the transition of parenchyma cells toward specific immune-activated states.

To confirm the occurrence of *Fgr*-specific strategy in parenchyma cells, beyond the anatomical outer sheath, we examined the spatial expression patterns of trajectory-associated genes using RNA *in situ* hybridization. The gene encoding a secretory peroxidase, which was upregulated in the *Fgr*-specific strategy (Figure 6F), was selected as an indicator of the cell state transition events. RNA *in situ* hybridization revealed specific expression of this gene in the outer sheath of mock-treated coleoptiles at 1 DPI, with subsequent expression in parenchyma cells following infection by the adapted pathogen *Fgr* (Figure 6G). This gene’s expression was also detected in some parenchymal cells in coleoptiles challenged by the non-adapted pathogen *Foc* (Figure 6G). Together with the observed preference of *Fgr* for invading parenchyma (Figure 1F), these findings suggest the occurrence of *Fgr*-specific strategy within parenchyma cells.

### Adapted fungal pathogen *F. graminearum* induced the chlorenchyma cells into a state of low transcriptional activity in the early infection stage

The chlorenchyma is composed of cells that contain chloroplasts and have a pronounced capacity for responding to pathogen infections (Figure 4D). However, the chlorenchyma displays notable heterogeneity, with the development of chloroplasts varying depending on the cell’s position within the coleoptile (O’Brien and Thimann, 1965). This heterogeneity was demonstrated in our scRNA-seq data, where the expression activity of three photosynthetic components varied among the chlorenchyma cells (Supplementary Figure 9A). Upon projecting the total mRNA count of each cell onto UMAP embeddings, it was observed that chlorenchyma cells exhibited a variable level of total mRNA (Supplementary Figure 9B). Among the four subpopulations of the chlorenchyma cells, subpopulation 6 exhibited a lower total mRNA level than other cells (Supplementary Figure 9B and 9C), indicating a low transcriptional activity. Further differential abundance analyses between fungal treatments and the mock at 1 DPI revealed that chlorenchyma cells from *Fgr* and *Foc* infected samples were enriched in opposing states with varying photosynthetic pathway activities (Supplementary Figure 9D). In addition, RNA velocity analysis suggests that the chlorenchyma cells undergo active transcriptome reprogramming in both mock and fungal treatments (Supplementary Figure 7B). These findings underscore the cellular dynamics of chlorenchyma in response to different fungal infections.

To delineate these transcriptional events, we conducted a pseudotime analysis of the chlorenchyma cells. A bifurcated trajectory was fitted in the transcriptional space of the chlorenchyma, depicting cells progressing toward two opposing states (Figure 7A). One cell state was characterized by high photosynthetic pathway activity (Supplementary Figure 9A), while the other exhibited lower total mRNA levels and reduced photosynthetic pathway activity (Supplementary Figure 9A and 9B). As revealed by the abundance analysis, cells exposed to different pathogens were predominantly enriched in these two transcriptional states (Figure 7B and Supplementary Figure 9D). Thus, branch 2 of the trajectory represents chlorenchyma progression towards the *Foc*-specific state of high photosynthetic pathway activity, whereas branch 1 depicts the progression to the *Fgr*-specific state of low transcriptional activity (Figure 7A and 7B).

**Figure 7.**
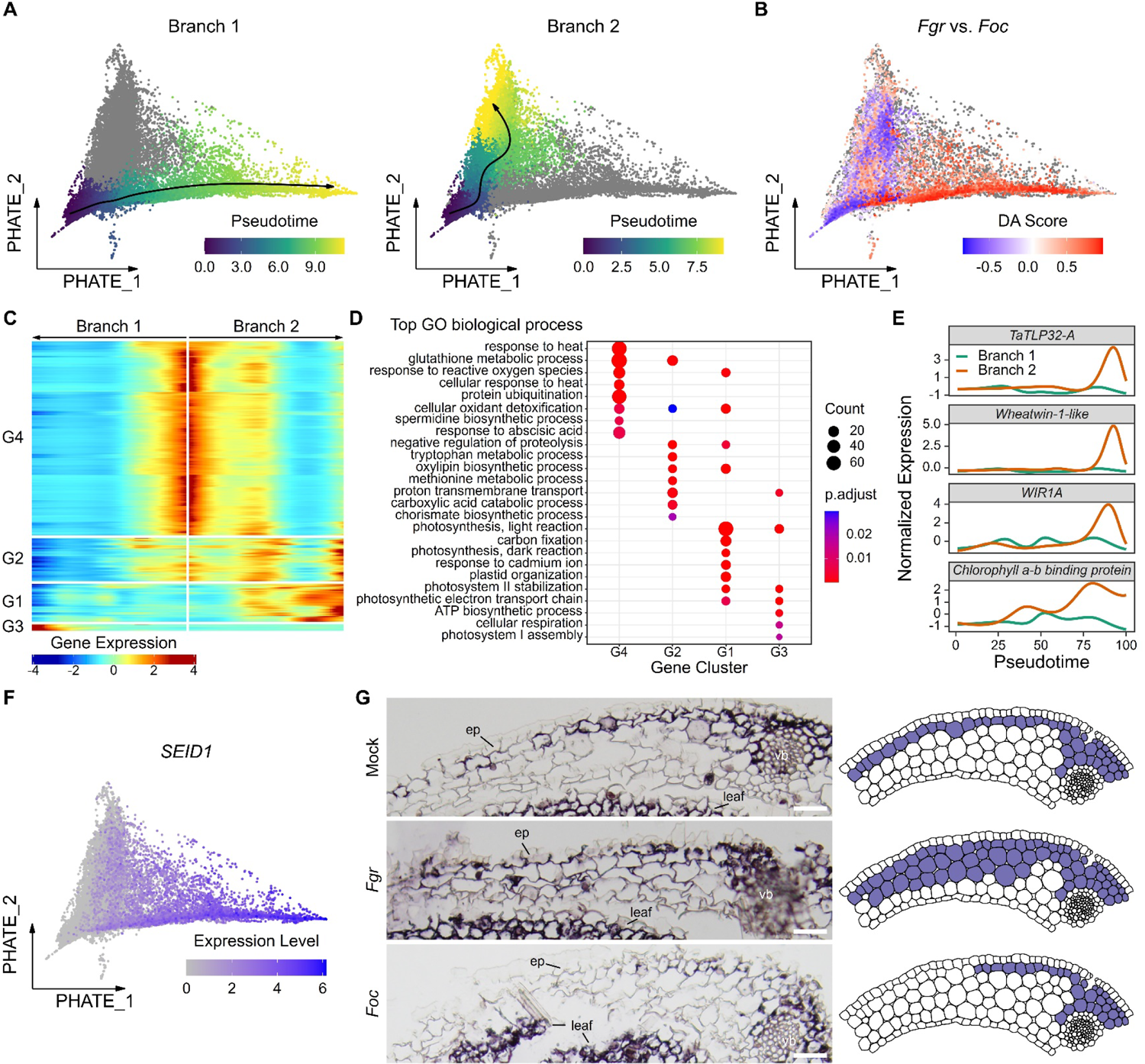
Chloroplast-containing cells transitioned to opposing transcriptional states upon the invasion of two *Fusarium* species (A) The trajectories inferred within the chlorenchyma (chloroplast-containing cells) depict a transcriptional switch in response to fungal invasion. The fitted principal curves were illustrated with arrows in black. Each dot in the PHATE projection, representing a cell, is color-coded according to its pseudotime value. (B) Differential abundance analysis reveals two chlorenchyma states, enriched in cells exposed to adapted and non-adapted pathogens. Cells in the PHATE projection are colored based on the differential abundance score (DA Score). Large DA Scores (red) indicate a higher abundance of cells from the *Fgr*-infected samples, while smaller scores (blue) are associated with the *Foc*-infected samples. Grey dots represent cells not classified into *Fgr* or *Foc* groups. (C) A heatmap illustrates the expression patterns of genes differentially expressed along pseudotime in two divergent trajectories. Arrows mark the trajectory directions, with genes categorized into four clusters by k-means clustering based on expression patterns. (D) Gene Ontology enrichment analysis of the gene modules from (C) characterizes distinct transcriptional states of the chlorenchyma cells in response to adapted and non-adapted *Fusarium* species. (E) The defense and photosynthetic genes were upregulated in the progression of the chlorenchyma to the *Foc*-specific state. (F) The *Subepidermal Intrinsic Disorder 1* (*SEID1*) gene was prominently expressed in a subpopulation enriched in cells from samples inoculated with the adapted pathogen *Fgr*. (G) RNA *in situ* hybridization for *SEID1* confirms transcriptional state transitions in the chlorenchyma cells at 1 DPI. Anti-sense probes for the marker gene were hybridized under uniform experimental and imaging conditions. The right panel’s illustration depicts the expression domain of *SEID1* (left panel) under various conditions. Scale bars = 50 μm.

To characterize the transition of the chlorenchyma to these opposing transcriptional states, we identified four gene modules whose expression changes significantly in association with pseudotime (Figure 7C). Gene Ontology enrichment analyses revealed that immunity-related pathways, such as responses to heat, ROS, and abscisic acid, as well as the metabolism of tryptophan, oxylipin, methionine, and chorismate, enriched in G4 or G2 modules, were rapidly downregulated during the progression to the *Fgr*-specific state (Figure 7C and 7D). These pathways were, however, gradually downregulated or even upregulated in the progression towards the *Foc*-specific state (Figure 7C and 7D). Notably, module G1, enriched with response to ROS, oxylipin biosynthesis, and photosynthesis, showed an upregulation trend in the *Foc*-specific state progression, while module G3, enriched with ATP biosynthesis and cellular respiration pathways, exhibited late upregulation in the *Fgr*-specific state (Figure 7C and 7D).

In particular, the expression of defense genes increased during the transition to the *Foc*-specific state (branch 2), illustrated by the upregulation of genes such as *TaTLP32-A*, which encodes a thaumatin-like protein, the gene encoding Wheatwin-1-like (a PR4 protein) (Caruso et al., 1999), and the defense gene *WIR1A* (Bull et al., 1992) (Figure 7E). This trend was also observed in photosynthetic genes, including one encoding a chlorophyll a-b binding protein (Figure 7E). Collectively, the gene expression pattern associated with pseudotime suggests that photosynthesis was maintained, and immunity was activated during the chlorenchyma’s transition to the *Foc*-specific state. In contrast, the rapid reduction of photosynthetic pathway genes and the lack of induction of defense genes are features of the *Fgr*-specific state.

Among the 141 genes in the G3 module upregulated in the progression to the *Fgr*-specific state (branch 1), 36 encode highly similar, yet uncharacterized proteins, each sharing over 50% sequence identity (Supplementary Figure 10A and 10C). The wheat genome contains 40 members of this protein family, averaging 121 amino acids in length (Supplementary Table 7). Protein sequence searches in the MobiDB database (Necci et al., 2017) indicated they all possess an intrinsic disorder region of approximately 30 amino acid residues at C-terminus (Supplementary Figure 10C). RNA *in situ* hybridization assays revealed that four genes of this protein family were expressed in chlorenchyma cells adjacent to the outer epidermis of coleoptiles (Figure 7G and Supplementary Figure 10D). Based on their expression patterns, we named this family the Subepidermal Intrinsic Disorder (SEID) proteins (Supplementary Figure 10B and 10D). In conclusion, the expression pattern of the SEID protein family is a defining characteristic of the chlorenchyma’s transition to the *Fgr*-specific state of low transcriptional and photosynthetic activity.

To validate the transition of chlorenchyma to the *Fgr*-specific state upon infection, we performed RNA *in situ* hybridization assays using *SEID1* in coleoptiles at 1 DPI. The selection of 1 DPI is based on the time point at which the difference in cell abundance between the two opposing states is most pronounced (Supplementary Figure 9E). In mock treatments, *SEID1* expression was limited to the chlorenchyma cells adjacent to the outer epidermis (Figure 7G). However, as an indicator of the *Fgr*-specific state, *SEID1* expression expanded to more chlorenchyma cells in coleoptiles infected with the adapted pathogen *Fgr* (Figure 7F and 7G). Conversely, under non-adapted *Foc* infection, *SEID1* expression diminished (Figure 7G). These results validate that the adapted *Fgr* can induce a special state in the chlorenchyma cells during the early infection stage.

## Discussion

### Compartmentalization of plant immune responses

Previous understanding of the molecular interactions between plants and pathogens has focused on cell-autonomous responses, with little knowledge of how different cells coordinate to mount an effective defense. Our comparative study of wheat cell transcriptomic changes has revealed that the host response can be divided into distinct stages, with different cell types participating. This results in either resistance or susceptibility to non-adapted and adapted *Fusarium*, respectively. In response to non-adapted *Fusarium*, the wheat coleoptile deploys a cell-type-specific first wave of defense during 1 to 2 days post-inoculation (DPI). By 3 DPI, gene expression reverts to a state similar to that observed in the mock treatment, with a reduction in the expression of many defense genes, indicating the establishment of nonhost resistance.

During the first wave of defense against non-adapted *Fusarium* in wheat coleoptiles, multiple cell types collaborate to orchestrate immune responses. Different cell types exhibit varying abilities to sense pathogens, and the transcriptional regulation of PTI recognition and signaling does not uniformly occur within the same cell type. The epidermis and phloem exhibit more complete PTI pathways (various pattern recognition receptors, MLO-like calcium channels, putative calcium-binding proteins, putative NADPH oxidases, and WRKY transcription factors), but the outer sheath only enhances PTI recognition (e.g. WAK, Lec-RLK, and LRR-RLK receptors). To orchestrate immune responses throughout the coleoptile, communication between different cell types is required. As potential intercellular signal molecules, the upregulation of SA biosynthesis (e.g. PALs genes) is confined to the phloem and outer sheath, whereas NHP is mainly in the outer sheath (e.g. FMO-like and ALD-like genes). The upregulation of antifungal pathogenesis-related (PR) genes, such as PR-1, chitinases, ribonucleases, and thaumatin-like proteins, is also restricted to the phloem and outer sheath. However, the biosynthesis of several defensive secondary metabolites (e.g. lignin, isoflavones, HCAAs, and benzoxazinoids) involves different cell types other than the phloem and outer sheath. Our results indicate that the wheat’s immune response to non-adapted *Fusarium* is compartmentalized, with multiple cell types involved in the orchestration of immune recognition and signaling pathways.

The spatiotemporal regulation of plant immune responses, initially investigated in Arabidopsis roots, underscores the need for controlled immune activation to defend against pathogens while accommodating commensals (Tsai et al., 2023). It is now understood that plant immunity in roots is both compartmentalized and specialized, with cell identity playing a crucial role in the transcriptional reprogramming that leads to cell-specific responses to MAMPs (Chuberre et al., 2018; Zhou et al., 2020; Rich-Griffin et al., 2020; Tsai et al., 2023). While plant leaves are generally not in constant exposure to MAMPs and do not accommodate commensals as roots do, the necessity for controlled immune responses to balance development and defense remains critical. However, the spatiotemporal dynamics of leaf responses to pathogens remain largely unexplored. Recent advancements in scRNA-seq have facilitated a deeper understanding of the Arabidopsis leaf transcriptome following infection by various pathogens, shedding light on cell-type-specific immune responses (Nobori et al., 2023; Zhu et al., 2023a; Delannoy et al., 2023; Tang et al., 2023).

In wheat coleoptiles, a leaf-like organ, our research identifies a cell identity-constrained capacity to respond to *Fusarium* pathogens. In the pathosystem of Arabidopsis leaf infection by the fungal pathogen *Colletotrichum higginsianum*, a significant expression of Toll/Interleukin-1 receptor NLR (TNL) genes is noted in vasculature-related cells (Tang et al., 2023). However, this pattern is not observed in the wheat coleoptile upon infection with the non-adapted fungal pathogen *Foc*. Instead, distinct subsets of *CNL*s are upregulated in the epidermis and L2-derived mesophyll cells, respectively. The biosynthesis of the signaling molecule NHP, crucial for SAR activation, occurs in distinct cell types between Arabidopsis and wheat under biotic stress, with *ALD1* and *FMO1* genes involved in its synthesis showing enhanced expression in Arabidopsis phloem companion cells upon *Pseudomonas syringae* infection (Nobori et al., 2023). Conversely, in wheat, we observed significant upregulation of FMO-like and ALD-like genes mainly in the outer sheath upon *Fusarium* infection. Additionally, the compartmentalization of defensive secondary metabolite biosynthesis, such as camalexin and indole glucosinolate in Arabidopsis (Nobori et al., 2023; Delannoy et al., 2023; Tang et al., 2023), aligns with our observations on lignin, isoflavone, HCAAs, and benzoxazinoids biosynthesis in wheat coleoptiles. Collectively, our single-cell expression analysis reveals the compartmentalization of plant immunity and secondary metabolism within the coleoptile, thereby extending the understanding of the heterogeneity in plant immunity from roots to leaf-like organs.

### Pivotal role of phloem and outer sheath in wheat defense against *Fusarium* pathogens

The adapted *Fusarium* grows among all cell types at the wounding inoculation site mainly in an intercellular manner around 1 to 2 DPI, similar to non-adapted *Fusarium*. However, it invades more cells with a cell-to-cell mode around 3 DPI (Zhang et al., 2012; Jia et al., 2019), contrasting with the blockage of non-adapted *Fusarium* by the host. Our analyses demonstrate that the phloem and outer sheath possess the highest capacity for responding to *Fusarium* invasion. The immune activation in the phloem is more sustained and complete during infection, involving PTI recognition and signaling (MLO-like calcium channels, putative calcium-binding proteins, and numerous LRR-RLKs), several antifungal PR genes (chitinases and thaumatin-like proteins), many phenylpropanoid metabolism genes (PALs, C4Hs, and 4CLs), lignin biosynthesis, chitin catabolism, hydrogen peroxide catabolism, and cell wall biogenesis. Remarkably, the phloem’s responses to both adapted and non-adapted *Fusarium* species share similar biological processes during 2 to 3 DPI. This highlights phloem’s pivotal role in basal immunity beyond its physiological function of nutrient transportation. Nevertheless, in response to adapted *Fusarium*, the phloem’s responsiveness is suppressed at 1 DPI (e.g. MLO-like calcium channels and LRR-RLKs).

The response of the outer sheath to non-adapted *Fusarium* is most drastic in the coleoptile, characterized by increased expression of multiple pattern recognition receptors (WAK, Lec-RLK, and LRR-RLK), antifungal PR genes (including PR1, chitinases, thaumatin-like, etc), numerous phenylpropanoid metabolism genes (PALs, C4Hs and 4CLs), putative SA biosynthesis genes, and a small portion of lignin biosynthetic genes. However, in response to adapted *Fusarium*, the coleoptile defense is delayed compared to non-adapted *Fusarium* in outer sheath cells, including lignin polymerization (i.e. Dirigent proteins), phenylpropanoid metabolism, tryptophan biosynthesis, suppression of photosynthesis, and suppression of hydrogen peroxide catabolism. Further single-cell level analysis revealed that the outer sheath cells display various immune responses in mock or fungal treatments. Therefore, the outer sheath may be naturally in a state of immune activation.

Notably, higher plants deploy border cells that perceive MAMPs and activate defenses, acting as protective “sentries” around the root meristem (Plancot et al., 2013; Chuberre et al., 2018). In Arabidopsis, defense mechanisms are restricted to peripheral root cap cells, preserving meristematic activity of root tips despite biotic stresses (Emonet et al., 2021). Additionally, the endodermis, representing the innermost cell layer of the cortex surrounding the stele, plays a pivotal role in root defense. It prevents systemic colonization by numerous fungal pathogens that typically invade roots radially before spreading via the vasculature to the foliage. Interestingly, a similar cell layer surrounding the vein is observed in stems and leaves (Martins et al., 2011), and is regulated by analogous mechanisms (Shaar-Moshe and Brady, 2023), although the terminology “endodermis” is controversial (Martins et al., 2011). The outer sheath in leaves, developed from the ground meristem like the root endodermis, serves a comparable cell type (Sakaguchi and Fukuda, 2008; Nelson, 2011; Martins et al., 2011; Zeng et al., 2016; Sedelnikova et al., 2018). Our scRNA-seq analysis of the wheat coleoptile revealed that the outer sheath is the most responsive cell type to pathogen infection and may have specialized immune functions. These findings imply that plants have evolved to confer greater immunity to certain cell types to protect vulnerable but vital tissues.

While the outer sheath is considered a specialized immune-active cell type protecting the vasculature of the coleoptile, these cells may represent a common transcriptional state of L2-derived mesophyll rather than being an anatomical cell type. The coleoptile, like roots, faces a complex soil microbiome and protects the tender shoot during seed germination. The “stomatal plugs” formed by soil-borne micro-organisms are extremely common, and these organisms are often found at some depth in the coleoptile tissues, usually in the intercellular airspaces (Thimann and O’Brien, 1965). The wheat coleoptiles used for this study were grown on wet paper towels, which is not an axenic condition, thus environmental microbes might trigger a defense response. Altogether, our study highlights the significance of exploring the function of specialized cell types in immunity from an evolutionary perspective.

### Understanding the susceptibility of wheat to *F. graminearum*

The understanding of specific wheat cell types manipulated by *F. graminearum* and the biological processes involved in such susceptible interactions remains limited. The responsiveness of the phloem and outer sheath in the first wave of nonhost resistance is suppressed at an early stage of the susceptible interaction with *F. graminearum*, as discussed above. This indicates that the adapted pathogen *F. graminearum* effectively evades plant immunity. Intriguingly, chlorenchyma cells enter a state of low transcriptional activity upon *F. graminearum* infection at 1 DPI, contrasting with the maintaining of high photosynthetic activity by the non-adapted *Fusarium*. Given the chloroplast’s role in integrating PTI signaling and producing phytohormone precursors (Nomura et al., 2012; Kachroo et al., 2021), the immune evasion of *Fgr* at the early stage may result from the “silencing” of chlorenchyma cells. This is reminiscent of the report indicating that, in the wheat spike, *F. graminearum* causes chlorenchyma cell death prior to direct contact, resulting in spike bleaching (Brown et al., 2010). Furthermore, the virulence factor fusaoctaxin A of *F. graminearum* can suppress the expression of chloroplast genes and disrupt the subcellular localization of chloroplasts in the wheat coleoptile (Jia et al., 2019). Therefore, these studies indicate that the virulence factors of *Fgr* may suppress the activity of the chlorenchyma cells.

In contrast to the compartmentalization and rapid onset of nonhost resistance, the adapted pathogen *F. graminearum* during 2 to 3 DPI elicits a more extensive, stronger, but delayed transcriptional activation of PTI, ETI, and SAR in multiple cell types, accompanied by the suppression of HCAA biosynthesis and lignin polymerization in outer sheath cells. The invasive growth of *F. graminearum* in the intercellular spaces of the parenchyma induces surrounding cells to transition into divergent immune-activated states in the later stages, differing from the limited nonhost resistance. However, the competition between immune and susceptible processes drives the transition of parenchyma cells towards a specific state that is favored by *F. graminearum*. The response of the wheat coleoptile reflects the infection strategy of *F. graminearum*, which is characterized by covert initial penetration and proliferation, followed by overt destruction (Zhang et al., 2012). Therefore, as a hemibiotrophic fungal pathogen, *F. graminearum* may manipulate the defense strategies of the parenchyma to induce susceptible processes and boost excessive immune responses to more cell types in the necrotrophic phase.

In conclusion, our study highlights the compartmentalization of plant immunity within the wheat coleoptile and identifies pivotal roles of the phloem and outer sheath. The mechanism of wheat’s susceptibility is revealed by the strategy of *Fgr* to escape plant immune responses by “silencing” the chlorenchyma in the early biotrophic phase and manipulating the defense strategies of the parenchyma during the necrotrophic phase. Therefore, this research provides a foundational resource to identify novel candidate genes for developing FHB-resistant wheat cultivars (Supplementary Table 2, 5, and 6), while minimizing growth penalties through precise expression of defense-related genes or silencing of susceptibility genes in targeted cell types. Additionally, our single-cell expression profiles offer a reference for *in silico* dissecting bulk RNA-seq data of the coleoptile under various conditions based on deconvolution methods (Newman et al., 2019; Chu et al., 2022), enabling the inference of cell-type-specific gene expression profiles without physical cell isolation. Despite the limitations of scRNA-seq, the advent of single-nucleus omics approaches in wheat (Zhang et al., 2023) presents new opportunities for exploring the transcriptional landscape of the spike in response to *Fgr* infection, paving the way for further advancements in our understanding of plant immunity and disease resistance mechanisms.

### Limitations

The requirement to generate protoplasts for plant scRNA-seq experiments introduces a challenge, as this process can induce transcriptional changes. However, excluding protoplast-inducible genes from single-cell datasets may lead to the loss of valuable information, especially for non-model plants. In this study, we employed mock controls at every time point and treatment to identify meaningful biological signals. Therefore, analyses of wheat cell responses under fungal treatment in comparison to mock treatment should be able to exclude gene expression changes caused by protoplasting alone, although it may miss those changes shared between immune responses and protoplasting responses.

Our findings derived from compositional analysis may be biased due to differential cell capture efficiency, influenced by the extent of cell wall enzymatic hydrolysis. Specifically, the proportion of coleoptile cells in our scRNA-seq data might be distorted by the absence of xylem cells. Moreover, the challenge of accurately defining cell types may complicate the interpretation of our analyses (Morris, 2019; Zeng, 2022). The distinction between cell states and cell types remains elusive, and pseudo-bulk expression analysis still suffers from Simpson’s Paradox.

## Methods

### Plant and Fungal Materials

The bread wheat (*Triticum aestivum L.*) cultivar Bobwhite was used for infection, scRNA-seq, and LCM-seq. The seeds were submerged in water, rinsed overnight, and then germinated on water-soaked paper towels at 25 °C with a 12-h photoperiod and greater than 90% relative humidity for three days before inoculation.

The transgenic strain of *Fusarium graminearum* (*Fgr*) PH-1 (NRRL 31084) expressing the ZsGreen driven by the promoter of *Aspergillus nidulans* oliC was constructed as the procedure described previously (Zhang et al., 2012). The transgenic strain of *Fusarium oxysporum* f. sp. *cubense* (*Foc*) tropical race 4 (NRRL 54006) expressing GFP protein was constructed previously (Jia et al., 2019).

### Coleoptile Infection with *Fusarium*

The conidial suspensions of *Fgr* were prepared to a final concentration of 1×10^6^ conidia/mL. The fresh conidia suspensions of *Foc* were produced by inoculating aliquots from −80 °C frozen microconidia stocks into 100 mL of potato dextrose broth (PDB) and cultured for three days at 180 rpm in an orbital incubator at 25 °C. The cultures were then filtered through a nylon membrane (approximately 38 μm in diameter) to separate mycelia and centrifuged for 5 minutes at 3,000 × g. Pellets containing fresh microconidia were resuspended in sterile water and adjusted to 1×10^6^ conidia/mL. Freshly prepared *Fgr* and *Foc* conidial suspension were inoculated to wheat seedlings according to the protocol (Jia et al., 2017).

### Preparation of Coleoptile Samples for scRNA-seq

Three days after wheat seed germination, the top 2 to 3 mm of coleoptiles were removed and 2.5 μL of conidial suspensions or sterile water was pipetted onto the wounds. Following inoculation, seedlings were grown in a growth chamber and coleoptiles were collected from the first to the third day post-inoculation (DPI). To investigate wheat’s transcriptional responses to pathogen *Fgr* and non-pathogen *Foc*, mock-inoculated (with sterile water) coleoptiles were used as controls. Two biological replicates of each treatment were collected at three time points (1 to 3 DPI). In addition, to measure the initial state of the transcriptome, coleoptiles with the top removed but otherwise untreated were immediately collected at 0 DPI. In conclusion, the experimental design included a total of ten sample types, each with two biological replicates (Figure 2A). Once the coleoptiles were harvested, they were first cut into 0.5-1 mm strips in a 0.6 M mannitol solution using fresh razor blades. The mannitol solution containing coleoptile strips was replaced with a freshly prepared enzyme solution (4% cellulase RS, 1% macerozyme R-10, 1% β-glucanase, 0.5% xylanase, 0.6 M mannitol, 10 mM MES [2-(*N*-morpholino)ethanesulfonic acid], 5 mM CaCl_2_ and 0.1% BSA [pH 5.7]) and then the coleoptile strips were digested for 3 hours at room temperature with 40-50 rpm shaking in dark. After digestion, the protoplast-containing enzyme solution was diluted with an equal volume of pre-chilled high osmolarity washing solution (1.2 M Glycerol, 10 mM MES, and 5 mM CaCl_2_). The protoplasts were subsequently filtered with a cell strainer (40 μm diameter, Falcon, Cat No 352340), centrifuged at 100 g and 4 °C for 10 min, and the pellet was resuspended in washing solution. After being on the ice for 15 minutes, the protoplasts were filtered through a nylon mesh with a pore size of 30 μm approximately in diameter and centrifuged again. To maintain proper osmotic pressure, the pellet was resuspended in a modified BD sample buffer supplemented with 1 M glycerol.

### scRNA-Seq Library Construction and Sequencing

The scRNA-Seq sequencing library was prepared according to the manufacturer’s instructions (Doc ID: 214062 Rev. 2.0, https://scomix.bd.com/hc/en-us/articles/360023044392-Instrument-Guides) for the BD Rhapsody system. Briefly, protoplasts were stained with fluorescein diacetate (Yeasen, Cat No: 40720ES03) and manually counted in the BD Scanner using a hemocytometer. The protoplast concentration was adjusted to 300-600 cells/μL. scRNA-seq libraries were generated by NovelBioinformatics Ltd., Co. (Shanghai) using the manufacturer’s reagents and instruments, and then sequenced on Illumina NovaSeq with 150 bp paired-end reads.

### Coleoptile Sectioning for LCM-seq

The top 4 mm of coleoptiles from three-day-old wheat seedlings were fixed and paraffin-embedded following the method described by (Zhang et al., 2018), and 10 μm thick cross-sections were obtained using a rotary microtome (Leica RM2235; Leica Biosystems Nussloch). To evenly mount the paraffin ribbons, they were floated on a drop of RNase-free water, and then the slide was dried on a 42 °C slide warmer. The target cells were isolated and captured using the ZEISS PALM Microbeam laser microdissection system.

### LCM-seq Library Construction and Sequencing

Total RNA was extracted from laser captured samples with the PicoPure RNA isolation kit (Applied Biosystems, Cat No./ID: 12204), yielding 3-7 ng per sample. The concentration and integrity of RNA samples were measured using Agilent Bioanalyzer 2100 (Agilent Technologies) as previously described (Tang et al., 2006; Tang et al., 2010). Following that, approximately 10 μg of antisense RNA was generated from each total RNA sample through 2-round amplification using the Arcturus RiboAmp HS PLUS Kit (Arcturus, Cat No./ID: KIT0525). The strand-specific libraries were then generated using the dUTP method and sequenced on Illumina NovaSeq with 150 bp paired-end reads.

### Bioinformatics Data Analysis of Single-Cell RNA-seq

#### Pre-processing of Raw scRNA-Seq Data

The raw scRNA-seq data were processed according to BD Rhapsody whole transcriptome analysis (WTA) pipeline v1.8 (Doc ID: 47383 Rev. 9.0 and Doc ID: 54169 Rev. 8.0, https://scomix.bd.com/hc/en-us/articles/360019763251-Bioinformatics-Guides). Briefly, nuclear genome sequences and annotations of hexaploid bread wheat cultivar ‘Chinese Spring’ were downloaded from Ensembl Plants (release 48, https://plants.ensembl.org/Triticum_aestivum), organellar genome sequences and annotations were downloaded from NCBI GenBank (MH051715 and MH051716), and then the genome index was built using STAR (v2.5.2b) (Dobin et al., 2013). Following the preparation of genome data, the pipeline was executed locally on a High Performance Computing cluster. The primary output of the BD Rhapsody WTA pipeline is a unique molecular identifier (UMI) count matrix where each row is a gene, and each column represents a cell. To remove low-quality cells, those with gene numbers over 3 median absolute deviations below the median (calculated on the log2-scale) were excluded. The cells were filtered for those having mitochondrial sequence lower than 5% and chloroplast sequence lower than 10%. The cells with low complexity where the log10-transformed number of genes per UMI was less than 0.8 were further discarded. In addition to quality control of the cells, genes with expression in fewer than 10 cells were also removed to reduce the size of the UMI count matrix. Since mRNA was captured in the half-open environment of BD Rhapsody, ambient RNA contamination was removed using SoupX (version 1.5.2) (Young and Behjati, 2020).

#### Integration of Multi-Sample Data

As cell divisions occur early in growth of the cereal coleoptile (Liptay and Davidson, 1972; Wiedenroth et al., 1990; Gibeaut et al., 2005), cell types are perceivably conserved in mock-treated samples from 0 DPI to 3 DPI, corresponding to 3 to 6 days after seed germination. We first integrated 8 datasets of mock-treated samples. After reviewing various methods have been developed to integrate scRNA-seq data from multiple samples, conditions, or laboratories, taking into account the ranking of preferred algorithms based on benchmarking studies (Chazarra-Gil et al., 2021; Luecken et al., 2022), we chose to test five popular algorithms: Seurat RPCA (Hao et al., 2021), Harmony (version 0.1) (Korsunsky et al., 2019), Scanorama (version 1.7.2) (Hie et al., 2019), batchelor FastMNN (version 1.12.3) (Haghverdi et al., 2018), and scVI (version 0.17.1) (Gayoso et al., 2022). According to the results of cell embeddings, clustering, and annotation, the Harmony algorithm was selected for its clear separation of cell clusters, well mixture of the samples, and a reasonable proportion of cell types. Before integrating Mock samples, Seurat (version 4.0.5) (Hao et al., 2021) was used to reduce the dimensionality of the datasets. In brief, the merged count matrix was normalized using the log-normalize method. The ‘vst’ method was used to identify the top 2,000 highly variable genes (HVGs), after which the normalized counts were scaled and centered based on the HVGs. To reduce the dimensionality of the scaled data, principal component analysis (PCA) was performed, and 20 PCs were used for integration. To correct batch effects of Mock samples, Harmony was run with parameter ‘theta’ set to 4 and 2 for the ‘sample’ and ‘time’ variables. Then, all the 20 datasets of samples with Mock, *Fgr,* and *Foc* treatments were integrated like that of Mock samples except that 30 PCs were used to run Harmony, and ‘theta’ was set to 4, 1, and 2 for the variables ‘time,’ ‘treatment,’ and ‘sample’, respectively.

#### Dimensionality Reduction, Clustering, and Cell Annotation

The dimensionality reduction for data summarization was performed in the previous batch-effect correcting step, further dimensionality reduction for visualization was based on the cell embeddings produced by the Harmony algorithm. The uniform manifold approximation and projection (UMAP) and t-distributed stochastic neighbor embedding (t-SNE) low-dimensional visualizations were performed by RunUMAP and RunTSNE functions of the Seurat toolkit (Hao et al., 2021). The FindClusters function of Seurat was used to identify cell clusters with a resolution of 0.6 and default parameters. To assign cell clusters to known cell types, we evaluated 1) the expression of known cell-type-specific marker genes in wheat coleoptile identified from the literature, 2) the expression of wheat genes homologous to known marker genes widely used in scRNA-seq of *Arabidopsis thaliana* leaves (Supplementary Table 1), and 3) the estimated abundance of different cell clusters in LCM-seq samples. The average normalized expression for the selected marker genes in each cell was computed and used to assign cell clusters to their putative cell types. For cell cluster proportion-guided assignments, the LCM-seq data of different niches in coleoptile were deconvolved with the CIBERSORTx algorithm (Newman et al., 2019), and the estimated proportions were then used to recover the spatial information of cell clusters defined by scRNA-seq.

After annotating the Mock dataset, cell cluster labels were transferred to fungi-inoculated samples based on a set of anchors establishing correspondences between cells of the reference and query datasets (Stuart et al., 2019). These labels were then used to guide the annotation of cells in the integrated dataset of all samples. The transcriptomes of cells from Mock, *Fgr*, and *Foc* treatments were integrated into a unified transcriptional space (Figure 4A), and all cells were grouped into 17 clusters via unsupervised clustering at the same resolution as the mock (Supplementary Figure 4A and 4B). As illustrated in the alluvial plot, the 17 cell clusters in the integrated dataset were further annotated to the same eight cell types derived from L1 to L3 as in the mock dataset by aligning the predicted cell cluster labels with those generated by unsupervised clustering (Figure 4B and Supplementary Figure 4C). To find out cell clusters impacted by fungal challenge, the distribution of cell label prediction scores in each cluster was illustrated in a violin plot (Supplementary Figure 4D), with lower scores indicating reduced prediction confidence (Stuart et al., 2019). Several clusters, including 1, 5, 9, 3, 7, 12, and 4, comprised a notable proportion of cells with low prediction scores, suggesting their scarcity in the mock dataset (Supplementary Figure 4D). This implies that these cell clusters emerge in response to fungal infection, underscoring the importance of annotating cells beforehand in mock datasets.

#### Aggregation of pseudo-bulk expression profiles

To evaluate the expression correlation among samples, the normalized and log-transformed counts of all cells within each sample were averaged. This average was employed to compute Pearson’s correlation. In the differential expression analysis between the treatment and control groups (i.e. mock-treatment) across each cell type, the raw counts from cells of the respective cell type were summed to form pseudo-bulk expression profiles. For the creation of expression heatmaps and the calculation of gene set activity, the mean of the normalized counts per million (CPM) for each cell type under various conditions was employed.

#### Cell Population Differential Abundance Analysis

A bootstrap resampling method implemented in scDC (Cao et al., 2019) was employed to capture the uncertainty associated with cell-type proportion estimates in replicated multi-sample scRNA-seq studies. A Bayesian model (scCODA) which accounted for multiple limitations of standard univariate statistical models was utilized to test compositional changes of each cell type between conditions (Büttner et al., 2021). The stomata (guard cell) was chosen as the reference cell type in the scCODA approach because it is mature and highly differentiated, less prone to changes caused by covariates. In all cluster-based compositional analyses described above, the granularity of the annotated cell types limits the resolution at which the transcriptional changes can be detected and interpreted. Hence, various cluster-free methods were used to reveal changes in subpopulations that were not discernible based on the employed cell type annotation. To characterize cell states that vary across samples and associate with sample attributes, i.e. days post-inoculation, we performed the co-varying neighborhood analysis (Reshef et al., 2022) based on the integrated Harmony (Korsunsky et al., 2019) transcriptional space. For differential abundance analysis between treatments with lower sample sizes, the DA-seq (Zhao et al., 2021) method was applied to the Harmony transcriptional space.

#### Gene Differential Expression Analysis

In the context of multi-sample, multi-condition single-cell differential expression analysis, pseudobulk-based methods have been considered the preferred approach in a benchmark study (Crowell et al., 2020), owing to their superior performance and stability. Consequently, DESeq2 (Love et al., 2014) was employed to identify cell-type-specific responses utilizing aggregated pseudo-bulk data. The Regulation Score used in heatmaps was calculated by summing the binarized log fold change of a gene in differential expression tests.

#### Cell Trajectory Inference

To assess the transcriptional dynamics of scRNA-seq data, RNA velocity was estimated using scVelo (Bergen et al., 2020) and UniTVelo (Gao et al., 2022). The expression matrices of spliced and unspliced mRNA in each sample were generated with the velocyto tool (La Manno et al., 2018), and then filtered and normalized using scVelo. The first and second order moments were computed among nearest neighbors in the Harmony space, and RNA velocity was then estimated by using the unified time model of UniTVelo. Cell population connectivity was estimated utilizing the partition-based graph abstraction (PAGA) method (Wolf et al., 2019). By integrating the single-cell data topology as elucidated by PAGA with RNA velocity analyses, we logically deduced potential transcriptional transition trajectories among cells. By incorporating this priori knowledge, trajectory inference was further refined using Slingshot (Street et al., 2018) which has been previously recognized as a top performer in tree-type trajectory analysis in a benchmarking study (Saelens et al., 2019). For the identification of differentially expressed genes along these trajectories, tradeSeq (Van den Berge et al., 2020) was employed.

#### Wheat Gene Pathway Analysis

The Gene Ontology (GO) annotations for the wheat genome were retrieved from the Ensembl Plants BioMart web server (release 52, https://plants.ensembl.org). Over-representation analysis and gene set enrichment analysis were conducted using the clusterProfiler software (Wu et al., 2021). To characterize cell-type-specific responses in wheat, GO enrichment analysis was performed on the differentially expressed genes. Due to enormous redundancy in the GO hierarchical graph, representative biological processes were manually selected based on semantic similarities among GO terms, though raw data are also provided (Supplementary Table 3). To assess pathway activity, the Gene Set Variation Analysis (GSVA) method (Hänzelmann et al., 2013) was utilized, employing pseudo-bulk Counts Per Million (CPM) expression profiles. Additionally, the AddModuleScore function of the Seurat package (Hao et al., 2021) was applied to single-cell expression matrices for pathway activity evaluation.

### Generation of Expression Matrix for LCM-seq

The sequencing data from LCM-seq were aligned to the genome index using the same alignment software as employed for scRNA-seq. Given that the mRNA in LCM-seq was amplified via *in vitro* transcription, resulting in antisense mRNA being utilized for library construction, the count matrix was produced using the featureCounts software (Liao et al., 2014) with the strand-specific read counting parameter set to “fr-secondstrand” to correctly account for the antisense orientation.

### RNA *in situ* hybridization

Candidate genes of wheat cultivar Bobwhite were cloned into pUC19. Antisense and sense RNA probes were synthesized *in vitro* using T7 RNA polymerase (Roche, Cat No./ID: 10881767001) and DIG RNA Labeling Mix (Roche, Cat No./ID: 11277073910). Tissues were fixed and embedded according to the procedure described in (Zhang et al., 2018), and the paraffin-embedded samples were sectioned (8 μm thick) with a rotary microtome (Leica RM2235; Leica Biosystems Nussloch). The slides were deparaffinized with xylene, rehydrated with a gradient of ethanol, and digested at 37 °C for 30 minutes with 1 mg/mL of proteinase K (Sigma, Cat No./ID: P6556). The slides were then dehydrated with a gradient of ethanol and hybridized with the probe overnight in a 52 °C oven. After sufficient washing, the non-specific sites were blocked with 1% Blocking Reagent (Roche, Cat No./ID: 11096176001). The slides were incubated with Anti-Digoxigenin-AP Fab fragment (Roche, Cat No./ID: 11093274910) for 90 minutes, followed by 1-3 days in an NBT/BCIP (Roche, Cat No./ID: 11681451001) solution. Images were obtained using an Olympus BX53 microscope.

### Histological Stains

For the observation of lignin deposition in wheat coleoptiles, the uppermost 10 mm segment of wheat coleoptiles was harvested, fixed, and subjected to staining with Basic Fuchsin, employing a ClearSee-based protocol (Kurihara et al., 2015; Ursache et al., 2018). Hand sections, with a thickness of 0.5 ∼ 0.8 mm, were prepared and imaged using a fluorescence microscope (OLYMPUS BX51) and a confocal microscope (OLYMPUS FV10C-W3).

## Data availability

The raw sequence data reported in this paper have been deposited in the Genome Sequence Archive in National Genomics Data Center, China National Center for Bioinformation / Beijing Institute of Genomics, Chinese Academy of Sciences that are publicly accessible at https://ngdc.cncb.ac.cn/gsa.

## Code availability

Vignettes for re-creating key analysis steps are available at https://github.com/altairwei/rhapsody-wta.

## Acknowledgments

We thank Dr. Zhihua Zhou and support from the Registry and database of bioparts for synthetic biology (https://www.biosino.org/npbiosys). This work was supported by Biological Breeding-Major Projects (2023ZD04070) and the Natural Science Foundation of China (32372479 to WT), the Strategic Priority Research Program of the Chinese Academy of Sciences (XDB27040212).

## Author Contributions

W.T. and W.W. conceived the project. W.W. performed data analyses and scRNA-seq experiments. S.L. performed LCM, RNA *in situ* hybridization, and microscopy. D.Z. participated in data analysis. W.T. and W.W. wrote the manuscript.

## Legend of Supplementary Figures

**Supplementary Figure 1.**
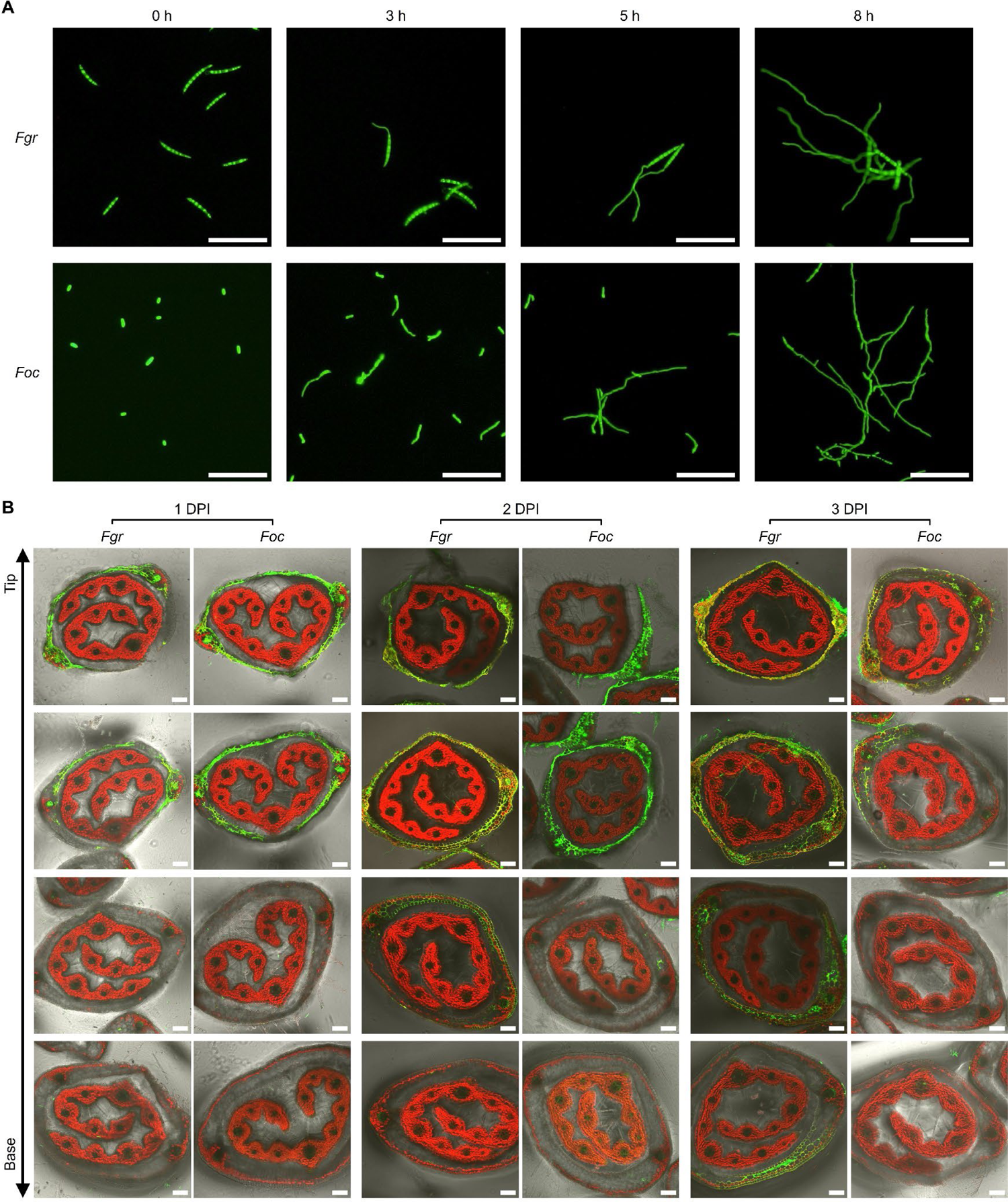
The *in vitro* germination of *Fgr* macroconidia and *Foc* microconidia, and their infection progression *in planta* (A) The germination process of the macroconidia of *Fgr* and microconidia of *Foc* in liquid Complete Medium. Scale bar = 100 μm. (B) Micrographs of serial hand cross-sections of coleoptiles infected by *Fusarium* pathogens reveal the invasion patterns and tissue preferences of the fungal pathogens. The fluorescence of GFP-carrying fungal hyphae (green) was merged with the auto-fluorescence of the wheat cell wall (green) and chlorophyll (red) in the same image. The serial cross-section images were arranged in order from the tip to the base, according to the distance from the inoculation sites. Scale bar = 100 μm.

**Supplementary Figure 2.**
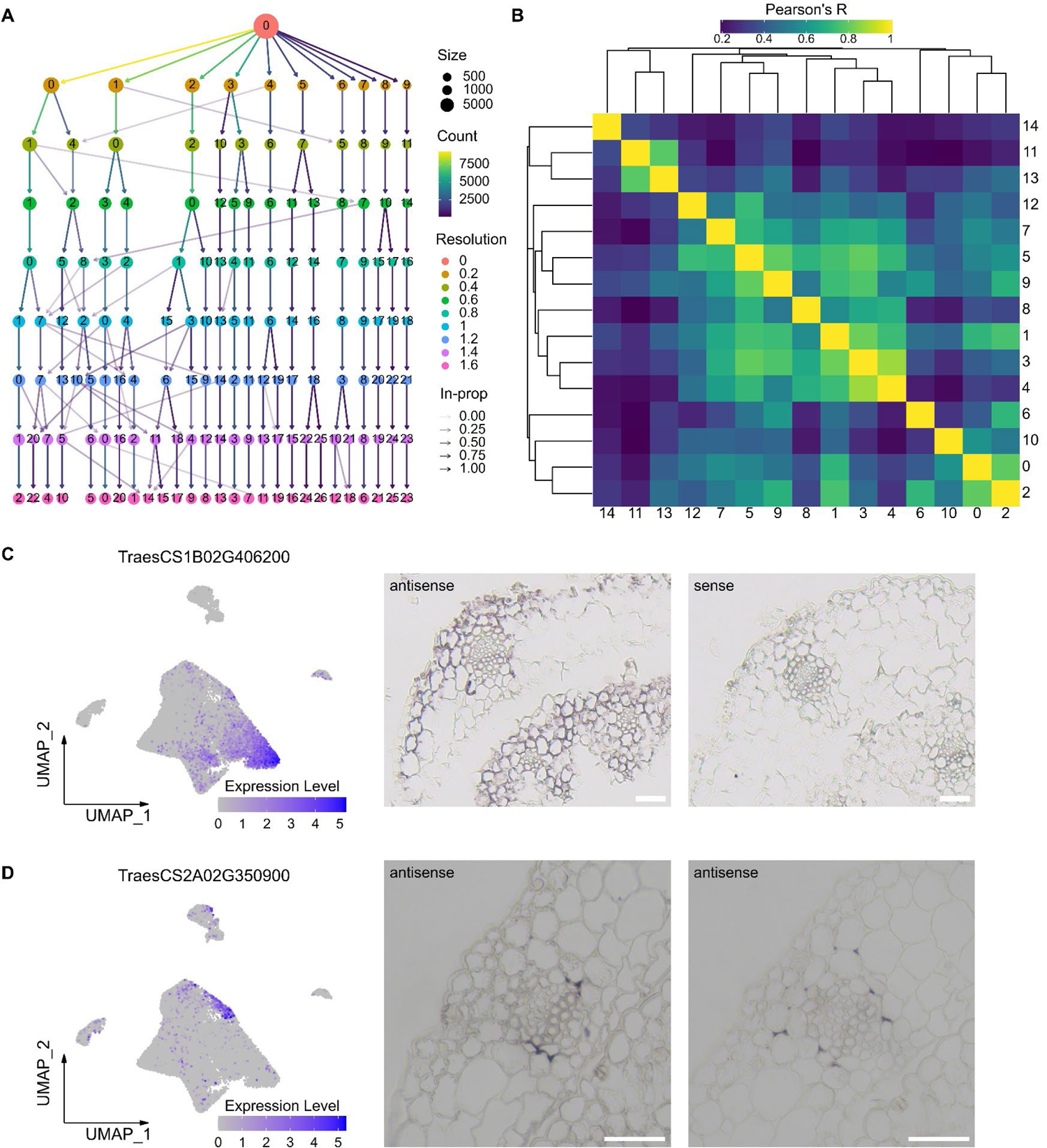
Supporting evidence for cell classification and annotation of Mock samples (A) The optimal resolution for grouping cells of Mock samples with similar expression profiles is guided by cell clustering trees. This plot illustrates the relationships between cell clusters at different resolutions and how cells move as the number of clusters increases. Size, cluster size. Count, the number of cells on the edge of the tree. In-prop, the ratio between the number of cells on the edge and the number of cells in the cluster it goes toward. (B) The heatmap shows the correlation of the average expression of 15 cell clusters, revealing their phylogenetic relationship. (C) RNA *in situ* hybridization of a gene specifically expressed in chloroplast-containing cells (left panel) validates the spatial localization of these cells (right panel). Scale bar = 50 μm. (D) RNA *in situ* hybridization of a gene expressed specifically in cell cluster 12 (left panel) showed that its transcripts accumulated in the outer sheath cells (right panel). Scale bar = 50 μm.

**Supplementary Figure 3.**
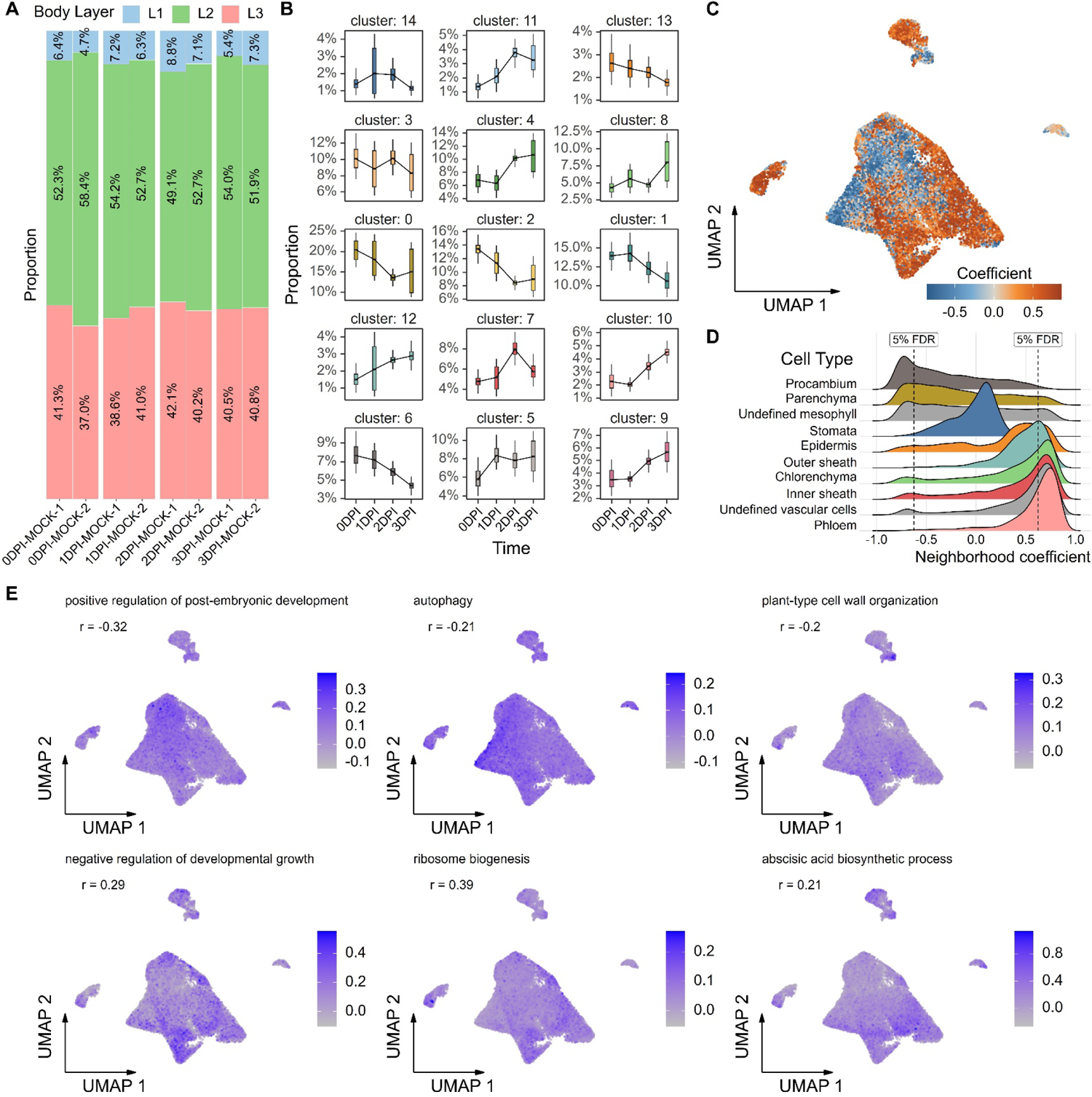
Developmental features of cell types elucidated the maturation and senescence of wheat coleoptile in the mock treatment (A) Average proportion of body layers remained relatively consistent over four days. (B) Changes in the proportion of cell clusters over time are displayed in the line chart. The uncertainty of proportion estimation is indicated by the box. (C) Co-varying neighborhood analysis identified cell states that vary across samples and associate with days post-inoculation. Each cell in UMAP embeddings is colored according to its neighborhood coefficient, with red values indicating a positive correlation with days post-inoculation, and blue values indicating a negative correlation. (D) The correlation of cell types with days post-inoculation is displayed by the distribution of neighborhood coefficients. The FDR < 0.05 thresholds are marked with vertical black dashed lines, highlighting that some cells from each cell type are included in the expanded (right) and the depleted (left) neighborhood over time. (E) The expression pattern of Gene Ontology modules that correlates positively or negatively with neighborhood coefficients reveals the developmental processes of the coleoptile. Pearson’s R is labeled at the top left.

**Supplementary Figure 4.**
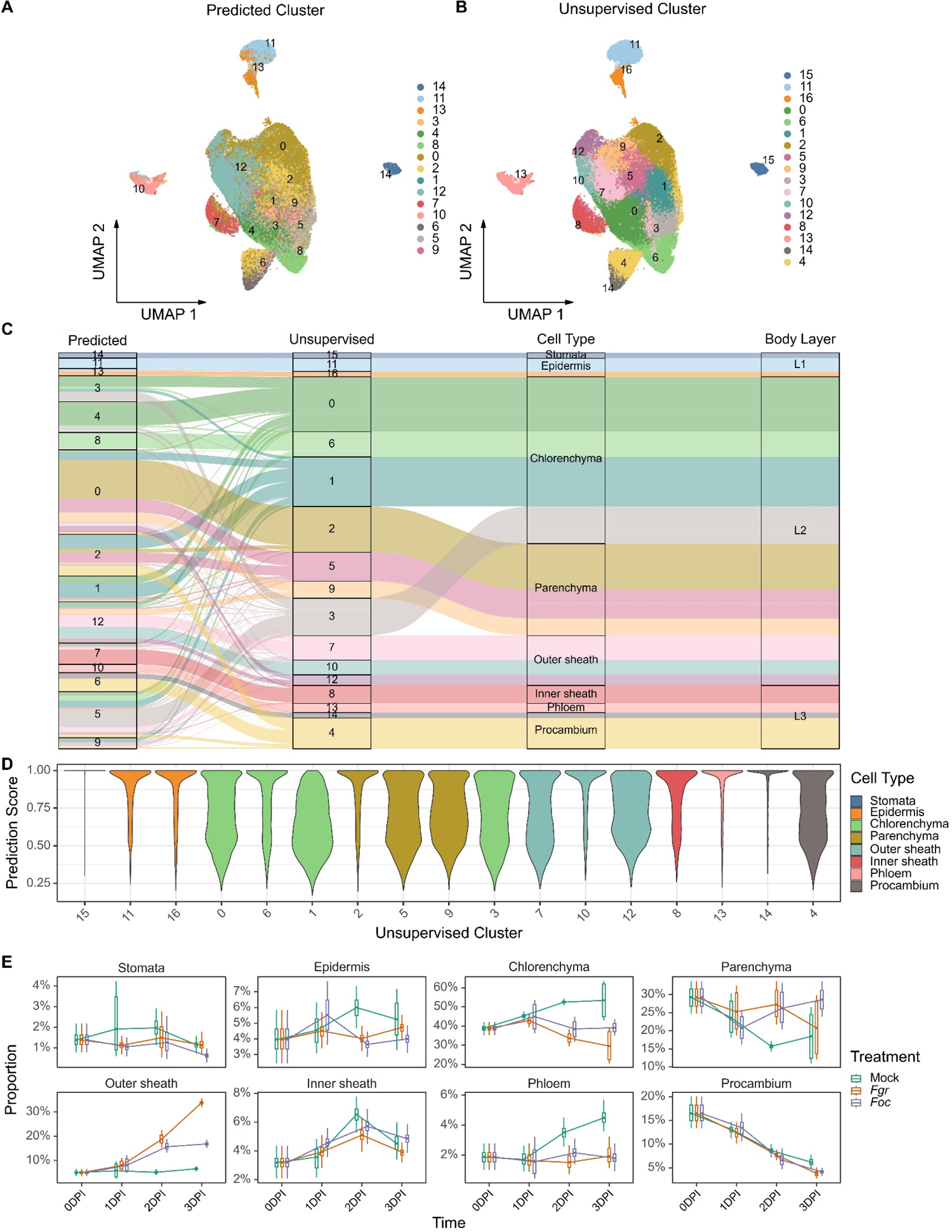
The prediction of cell types in pathogen-infected samples based on the annotated Mock dataset (A) 15 cell clusters of the integrated dataset were predicted from the Mock dataset. The UMAP projection displays cells as dots, which are colored according to their respective clusters. (B) Cells in the integrated dataset were grouped into 17 clusters using an unsupervised clustering method, not a prediction. (C) The alignment of the predicted cell cluster labels with those generated by unsupervised clustering identified 8 cell types of the integrated dataset. In the alluvial plot, cells are represented by lines that pass through different columns. (D) The distribution of prediction scores highlights cell clusters affected by fungal treatment. Lower scores indicate reduced prediction confidence, which suggests their likely absence in the mock dataset. (E) Changes in the proportion of cell types over time upon Mock, *Fgr*, and *Foc* treatment are displayed in the line chart. The uncertainty of proportion estimation is indicated by the box.

**Supplementary Figure 5.**
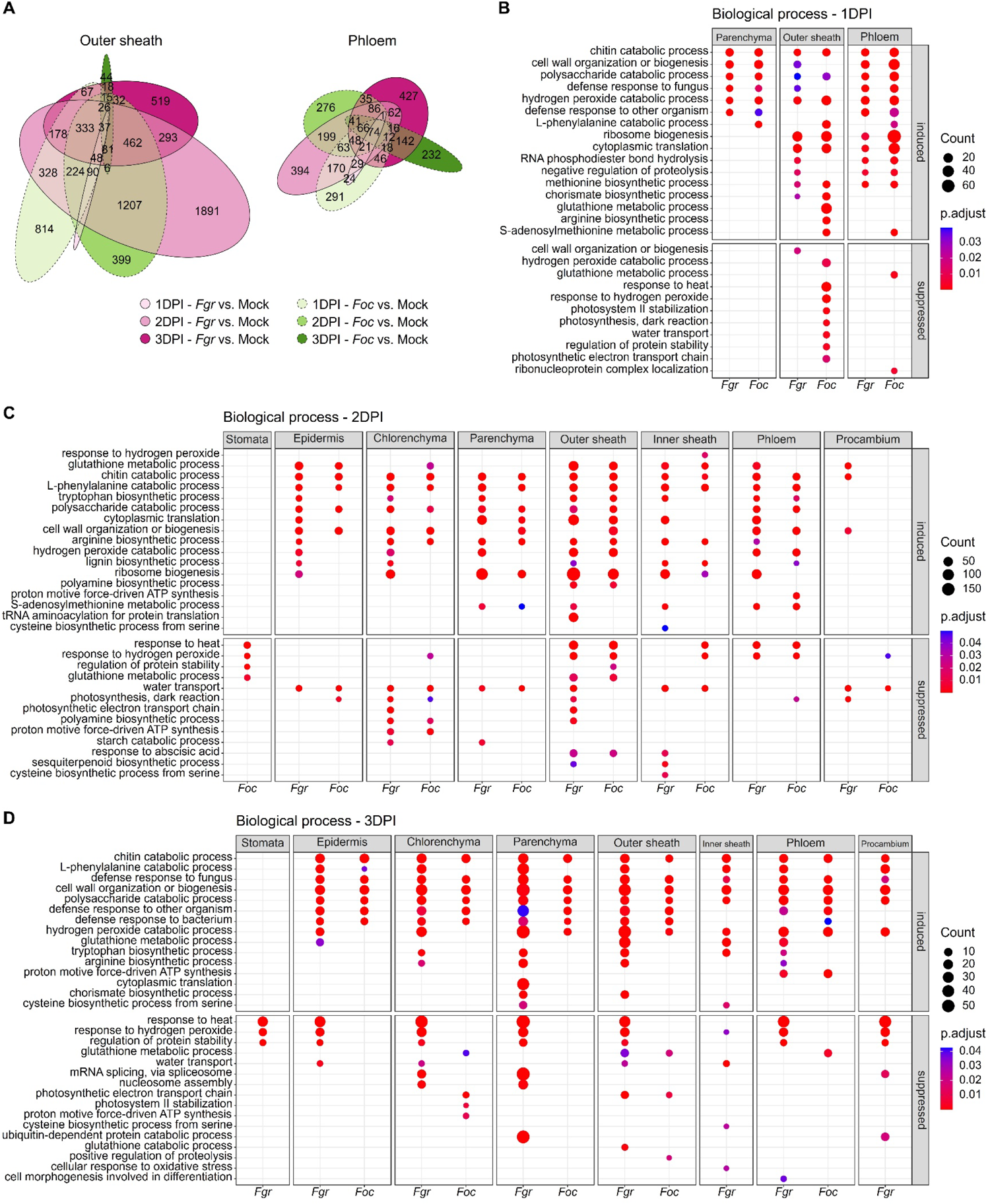
Top enriched GO biological processes in cell-type-specific responses (C) The area-proportional Euler diagrams of the DEG sets of the outer sheath and phloem indicate similarity among different times and treatments. Numbers in brackets specify the count of DEGs in each subset. (D) Simultaneous enrichment analysis was conducted on multiple sets of DEGs to identify commonly or specifically regulated biological processes in cell types of 1 DPI samples. The results were visualized as a dot plot with an x-axis representing fungal treatments, a horizontal facet panel representing different cell types, and a vertical facet panel representing DEG regulation type (induced or suppressed). The results of enrichment analysis for 2 DPI and 3 DPI are shown in (C) and (D), respectively.

**Supplementary Figure 6.**
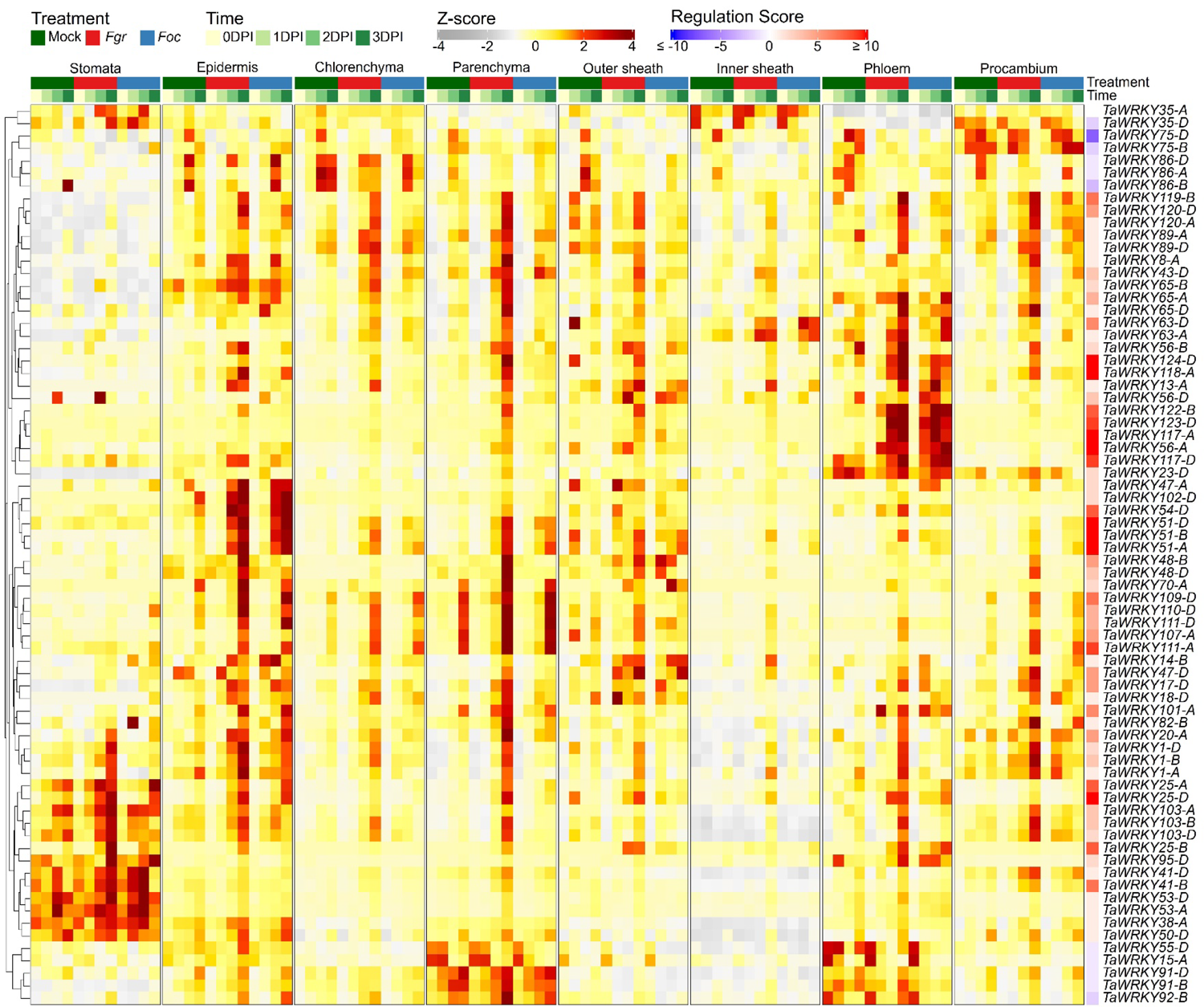
The adapted pathogen *Fgr* induced the extensive upregulation of WRKYs in most cell types The upregulation of WRKY transcription factors, which were restricted to the epidermis and phloem upon non-adapted pathogen *Foc* infection, was altered by the adapted pathogen *Fgr*. The heatmap displays the expression level (Z-score) of a single gene across eight cell types, with treatments shown from left to right within each cell type column. The treatments include mock-inoculated (green), adapted *Fgr*-inoculated (red), and non-adapted *Foc*-inoculated (blue). The heatmaps also show four distinct time points (0-, 1-, 2-, and 3-DPI) within each treatment block. The Regulation Score is displayed in a separate column to the left of the row names, using blue and red to indicate down- and up-regulation of genes in the coleoptiles.

**Supplementary Figure 7.**
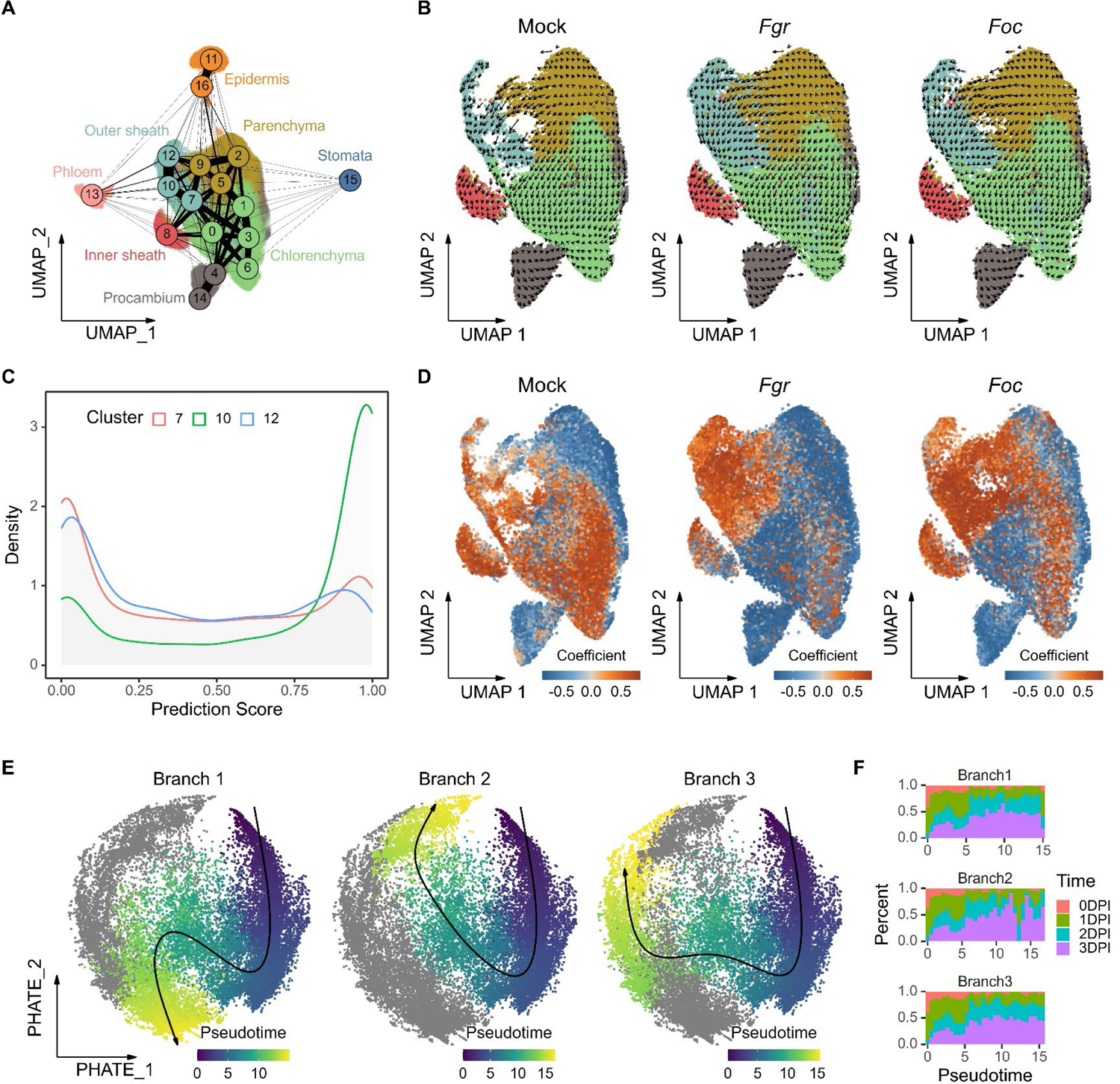
Identification and characterization of coleoptile’s spatiotemporal cellular interactions with *Fusarium* pathogens (A) Partition-based graph abstraction (PAGA) map quantifies the transcriptomic similarity of cell clusters. Different cell clusters may represent subpopulations of the same cell type, and therefore are coded with the same color. The PAGA map overlays the UMAP embeddings, where the nodes with the number identifier inside correspond to cell clusters, and edge weights quantify the connectivity between cell clusters. (B) RNA velocity vectors projected on UMAP embeddings reveal the future state of coleoptile cells. The speed and direction of the RNA velocity are indicated by arrows. Dots below the vector field in the UMAP embeddings represent cells and are colored by corresponding cell types. (C) The distribution of the prediction score, estimated when transferring cell types from Mock to infected samples, reveals varying confidence levels in subpopulations (cell clusters 7, 10, and 12) predicted as the anatomical outer sheath. (D) Co-varying neighborhood analysis identified cell states that vary across samples and associate with days post-inoculation. Each cell in UMAP embeddings is colored according to its neighborhood coefficient, with red values indicating a positive correlation with days post-inoculation, and blue values indicating a negative correlation. (E) Distinct trajectory branches fitted to parenchyma and outer sheath cells reflect unique interactions between the host and fungi. Each point in the PHATE embeddings, representing a cell, is colored according to the calculated pseudotime. (F) The histogram of inferred pseudotime for cells sampled from various days post-inoculation suggests asynchronous progression of cell state transitions.

**Supplementary Figure 8.**
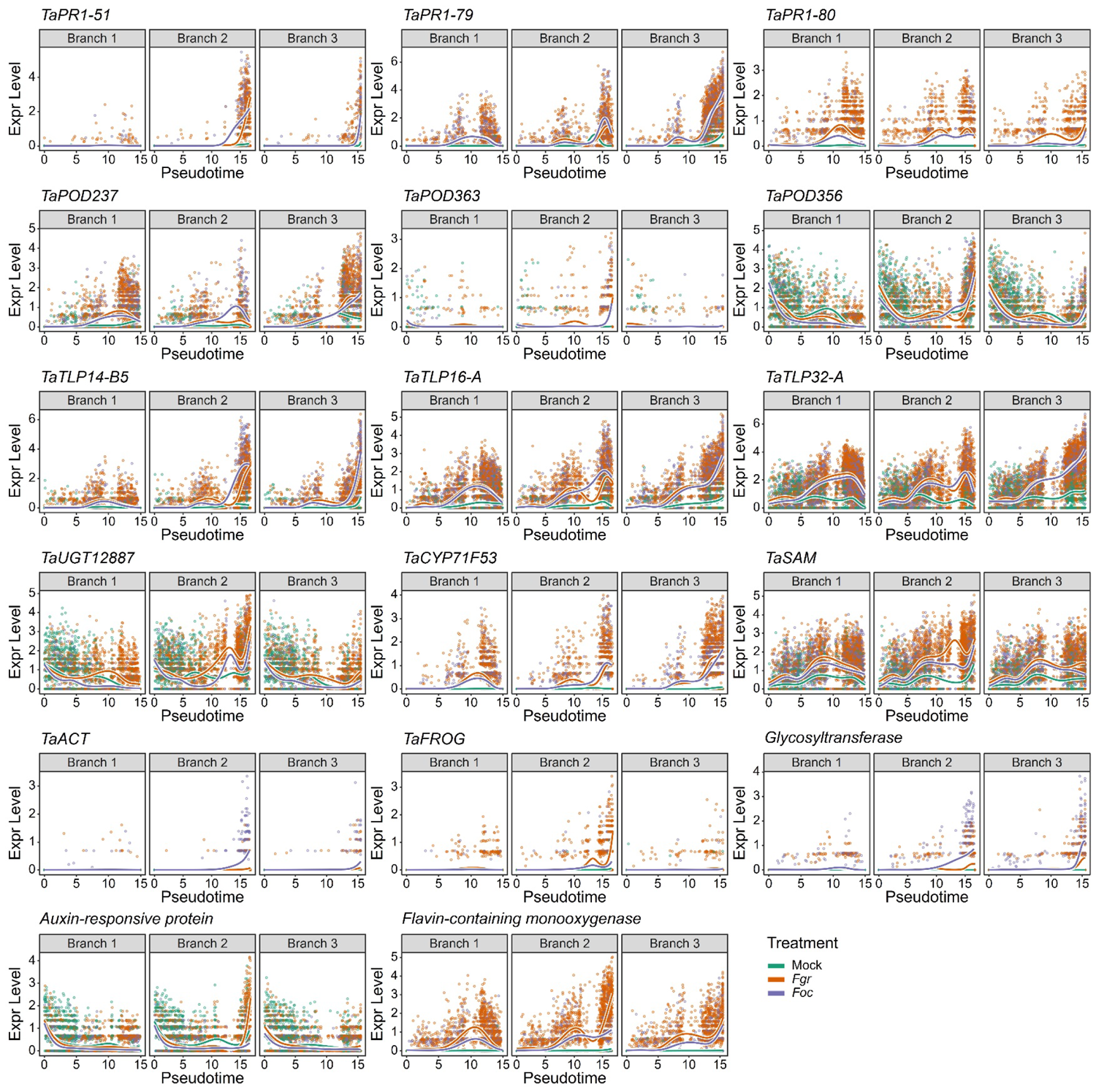
The expression pattern of immune genes along the trajectory depicts the parenchyma cells’ interaction with *Fusarium* pathogens The expression level of immune genes associated with the branches in the trajectory of the parenchyma cell state transition was compared between Mock, *Fgr,* and *Foc* treatments. Each dot represents a cell and is colored with its original treatments. Estimated expression smoothers for corresponding genes are shown as curves and colored with different treatments. *TaPR1*, *Triticum aestivum* pathogenesis-related protein-1. *TaPOD*, peroxidase. *TaTLP*, thaumatin-like protein. *TaUGT*, UDP-glycosyltransferase. *TaCYP71F53*, cytochrome P450 enzyme. *TaACT*, agmatine coumaroyl transferase. *TaSAM*, S-adenosyl methionine-dependent methyltransferase. *TaFROG*, *Triticum aestivum Fusarium* resistance orphan gene.

**Supplementary Figure 9.**
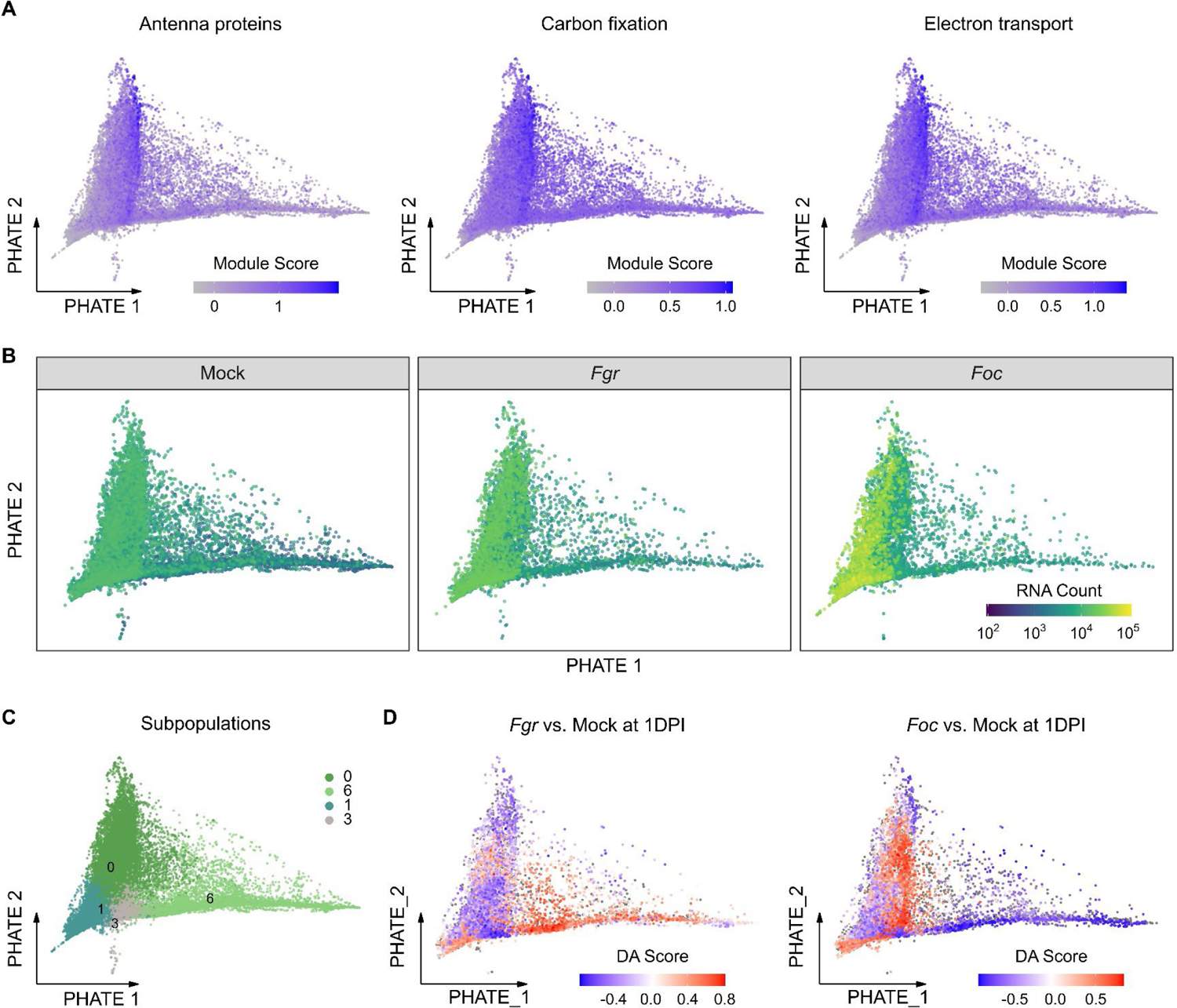
The chlorenchyma cell states were influenced by pathogen infection (A) The expression activity of three photosynthetic components varied among the chlorenchyma cells. The module scores summarize the expression of genes involved in photosynthesis antenna proteins, carbon fixation, and electron transport. These scores are then projected onto the PHATE embeddings of the chlorenchyma cells. (B) The total mRNA counts varied among the chlorenchyma cells, particularly between those belonging to two opposing states. The cells from mock-, *Fgr*-, and *Foc*-inoculated samples are displayed in each facet. (C) The chlorenchyma cells can be further clustered into four subpopulations with distinct transcriptional states. (D) Differential abundance analysis reveals that the chlorenchyma cells exposed to adapted and non-adapted *Fusarium* infection at 1 DPI were enriched in opposing states. Cells in the PHATE embeddings are colored based on the differential abundance score (DA Score).

**Supplementary Figure 10.**
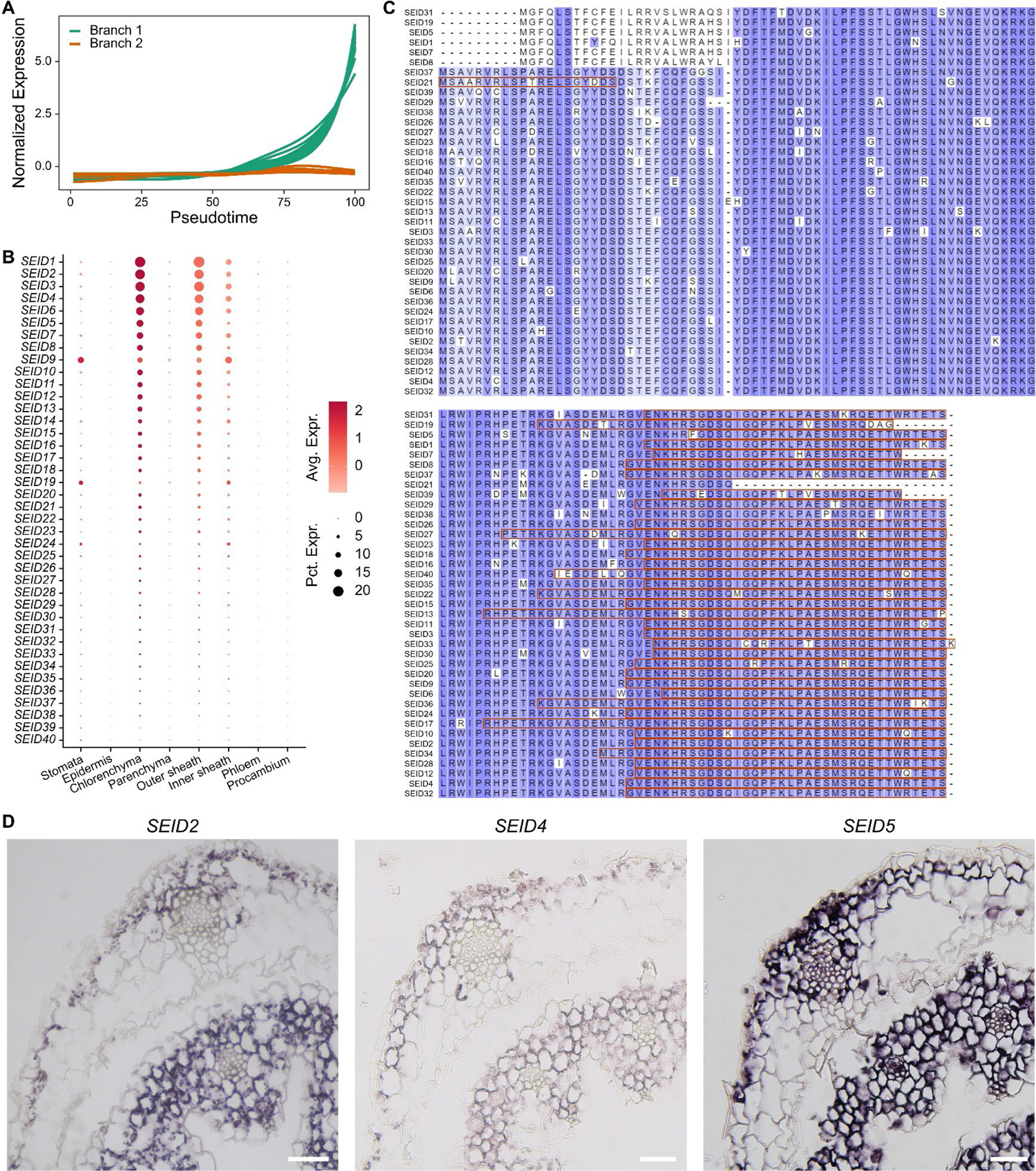
Identification of the Subepidermal Intrinsic Disorder (SEID) protein family as potential markers for the *Fgr*-specific state of the chlorenchyma (A) The expression of 36 *SEID* genes gradually increased as the chlorenchyma transitioned to the *Fgr*-specific state. The transition was characterized by two inferred trajectories, of which branch 1 leads to the *Fgr*-specific state. (B) The *SEID* genes were expressed primarily in the chlorenchyma of the coleoptile. The gene’s average expression (Avg. Expr.) and the percentage of expressed cells (Pct. Expr.) are indicated by the color and size of the dots, respectively. (C) The sequence alignment of SEID proteins demonstrates their conserved structure, with amino acid residues highlighted in blue to denote similarity. Brown boxes mark sequences as intrinsically disordered regions predicted by MobiDB-lite. (D) The subepidermal localization of *SEID* transcripts in the Mock-treated coleoptile at 1 DPI is displayed through RNA *in situ* hybridization. Scale bar = 50 μm.

## Supplementary Notes

**Supplementary Note 1. Expression analysis of well-annotated marker genes in the wheat coleoptile** We used a set of well-annotated marker genes with known expression patterns to indicate cell identity. The cluster 10 cells have the high and selective expression of the wheat sucrose importer *TaSUT1* (Figure 3B), which has been shown to specifically express in phloem companion cells of wheat leaves by RNA *in situ* hybridization and protein immunolocalization (Aoki et al., 2004). This suggests that cluster 10 cells belong to the phloem, in accordance with their spatial allocation as part of vascular tissue, as indicated by LCM-based RNA-seq deconvolution (Figure 3D). The cluster 6 cells exhibit high and specific expression of a wheat serine carboxypeptidase-encoding gene *CPIII* (Figure 3B), which has been demonstrated to be specifically expressed in early differentiating vascular cells (i.e. procambium) of wheat seedlings by *in situ* hybridization (Domínguez et al., 2002), indicating that the cluster 6 corresponds to the procambium. A wheat germin-like oxalate oxidase gene *gl-OXO*, whose expression predominantly accumulated in cluster 7 (Figure 3B), has been shown to express in sheath cells surrounding vascular tissue in wheat coleoptiles by RNA *in situ* hybridization (Caliskan and Cuming, 1998). This indicates that cluster 7 cells are vascular inner sheath in the coleoptile, which is also consistent with their spatial allocation of L3 derived from the LCM data. The clusters 11, 13, and 14 show high and selective expression of the Arabidopsis *High Carbon Dioxide* (*HIC*) homolog encoding 3-ketoacyl-CoA synthase 13 for the synthesis of long-chain fatty acids in cuticular wax, which is highly expressed in guard cells and adjacent epidermal cells (Liu et al., 2020), suggesting that the clusters 11, 13, and 14 are guard cells or epidermal cells. Cluster 14 is annotated to be guard cells based on the specific expression of homologs of an aluminum-activated malate transporter ALMT12 (Sasaki et al., 2010) and a guard cell-specific transcription factor AtMYB60 (Cominelli et al., 2011). Clusters 11 and 13 are annotated to be epidermal cells based on the specific expression of homologs of *Arabidopsis thaliana* epidermal marker genes: 3-KETOACYL-COA SYNTHASE encoding gene *FIDDLEHEAD* (*FDH*), homeobox protein-encoding gene *ATML1*, cutin biosynthetic acyltransferase encoding gene *DCR*, and chitinase *EP3* (Figure 3B and Supplementary Table 1), aligning with their spatial allocation (Figure 3D).

The mesophyll, constituting the largest proportion of the coleoptile, exhibits variation in morphology and function, but lacks a clear separation of different cell subpopulations. Two groups of mesophyll cells in the L2 layer are apparent in cross-sections of the coleoptile: greening and non-greening cells (Figure 1C). Homologs of *Arabidopsis thaliana* photosynthesis-related genes (*RBCS*, *CAB3*, *LHCB2.1*, and *AOC2*) were highly expressed in cell clusters 3, 4, 8, and 1 (Figure 3B). RNA *in situ* hybridization assays validated the location of these clusters in the L2 layer, showing distribution patterns similar to those of chloroplasts in the coleoptile (Figure 1C and Supplementary Figure 2C). These chloroplast-containing cells (clusters 3, 4, 8, and 1) were termed chlorenchyma. The non-greening cells in the L2 layer were grouped into cluster 0, constituting a substantial proportion (Supplementary Figure 3B), and were thus termed parenchyma. Notably, cluster 12, allocated to the vascular bundle, also expressed photosynthesis-related genes (Figure 3B and 3D), resembling the parenchymatous bundle sheath (outer sheath) in cereals, a layer of cells adjacent to the mestome sheath (inner sheath, cluster 7) (Williams et al., 1989; Zeng et al., 2016). RNA *in situ* hybridization of a gene specifically expressed in cluster 12 showed signals in bilateral parenchymatous cells surrounding the vascular bundle (Supplementary Figure 2D), confirming the identity of cluster 12 as the outer sheath. The remaining cell clusters with ambiguous identities, such as clusters 1, 2, 5, and 9, were assigned to either the undefined mesophyll or vasculature based on their expression similarity (Supplementary Figure 2B and Figure 3A). Therefore, the continuum of mesophyll cells in the L2 layer was categorized *in silico* into three cell types: chlorenchyma (clusters 3, 4, 8), parenchyma (cluster 0), and vascular outer sheath (cluster 12).

**Supplementary Note 2. The analysis of cell abundance and gene expression of the wheat coleoptile under mock treatment** Transcriptomic features of each cell type in mock samples mainly reflect cell growth and development states during coleoptile elongation, providing context for comprehending the coleoptile’s response to fungal invasion. In line with UMAP projection L1 and L3 cell clusters are isolated from L2 cell clusters, while mesophyll cell clusters within L2 (clusters 0-4, 8, 12) are very close to each other. We found that the proportions of the three body layers remained unchanged over four days (Supplementary Figure 3A), whereas the proportions of cell clusters within the L2 and L3 layers substantially changed (Supplementary Figure 3B). To understand the cellular transcriptomic dynamics during coleoptile development, co-varying neighborhood analysis (Reshef et al., 2022) was utilized to examine the correlation between cell abundance and growth time (0 to 3 DPI), as indicated by neighborhood coefficients. Projecting these coefficients onto UMAP embeddings revealed cells with positive or negative correlations with time scattered on either side of the transcriptional space (Supplementary Figure 3C). The results show that cell clusters 0, 1, 2, and 6 are younger (i.e. negatively correlated with time) and cell clusters 3, 4, 8, 12, and 7 are older (i.e. positively correlated with time) (Figure 3A and Supplementary Figure 3C). The density distribution of neighborhood coefficients across cell types reflected their temporal properties (Supplementary Figure 3D). Compared to stomata, which maintained consistent cell abundance, procambium, and parenchyma contained a higher proportion of younger cells, while chlorenchyma, vascular bundle sheath, and phloem comprised more mature cells. The trend for chlorenchyma cells to be older than parenchyma cells is consistent with the fact that coleoptiles are greener over time. Gene set enrichment analysis of genes with expression correlating with neighborhood coefficients unveiled the developmental processes underlying maturation and senescence (Supplementary Figure 3E). In summary, developmental features of the eight cell types elucidated the maturation and senescence of wheat coleoptile in the mock treatment.

**Supplementary Note 3. Expression pattern analysis of PTI, ETI, and SAR-related genes under fungal treatments** Concerning PTI immune recognition, a subset of genes encoding putative cell-surface immune receptors (PRRs), including three LRR-RLKs, two WAKs, and two Lec-RLKs, exhibited preferential upregulation upon infection with both *Fusarium* species from 1 to 3 DPI in the epidermis (Figure 5A). Distinct subsets of putative PRRs demonstrated specific upregulation: one comprising an LRR-RLK, two WAKs, and two Lec-RLKs in the outer sheath, and another, consisting of five LRR-RLKs, preferentially in the phloem (Figure 5A). This differential induction of non-overlapping PRRs across various cell types suggests that cells in the coleoptile do not have equal ability to sense pathogens.

In terms of PTI signaling components, putative MAPKs and calcium channels CNGC (MLO-likes) exhibited differential expression in response to *Fusarium* infection. Notably, in the phloem, 8 MLO-like genes showed significant upregulation, whereas 9 MAPK genes exhibited decreased expression upon infection with both *Fusarium* species (Figure 5A). Furthermore, the upregulation of three putative ROS producers (*TaNOX10*s), namely NADPH oxidase respiratory burst oxidase protein D homologs (RbohD), was consistent with the epidermis-specific induction of PRRs (Figure 5A). PTI signaling pathways can activate WRKY transcription factors, thereby enhancing the expression of defense-related genes (Ngou et al., 2022). Significantly, numerous WRKYs were upregulated upon non-adapted pathogen *Foc* infection, with distinct groups showing increased expression in the epidermis and the phloem (Supplementary Figure 6). The induction of PTI signaling components, including calcium channels, RBOHs, MAPKs, and WRKYs, varied among different cell types. Taken together, the epidermis and phloem exhibited roughly complete PTI pathways compared to the outer sheath, which only demonstrated upregulation of PTI recognition components (i.e. PRRs). These results suggest that the transcriptional regulation of PTI recognition and signaling does not uniformly occur within the same cell types.

Interestingly, the phloem-specific PRRs were upregulated as early as 1 DPI upon non-adapted pathogen *Foc* infection, whereas this upregulation was delayed until 2 DPI in the phloem during adapted pathogen *Fgr* infection (Figure 5A). Additionally, the subset of putative PRRs initially induced in the epidermis showed upregulation in the parenchyma and outer sheath by *Fgr* infection specifically at 3 DPI. At this juncture, 10 putative MAPK genes experienced increased expression across most cell types (Figure 5A and Supplementary Table 2). The cell-type-specific upregulation of WRKYs in nonhost resistance to *Foc* was also boosted by *Fgr* at 3 DPI, exhibiting more pronounced upregulation across most cell types (Supplementary Figure 6). Thus, the upregulation of PTI recognition and signaling components occurred at mismatched times and in inadequate cell types during the susceptible interaction with *Fgr*.

Nucleotide-binding leucine-rich repeat receptors (NLRs), crucial for ETI recognition and signaling, include 2151 putative *TaNLR*s carrying a nucleotide-binding domain in wheat (Andersen et al., 2020). However, only a small fraction, 23 (approximately 1% of 2151) *TaNLR*s, showed significant upregulation in at least one cell type upon *Fusarium* inoculation compared to mock-inoculated controls (Figure 5A and Supplementary Table 2). Among these, one *NLR*, lacking the LRR domain but carrying an NB-ARC domain, was specifically upregulated in the phloem, while others demonstrated significant increases across two to six cell types. In three L2-derived cell types, a group of *NLR*s, including three *CNL*s, one *NLR*, and one *NB*, were significantly upregulated upon non-adapted pathogen *Foc* infection. Conversely, a different group of *NLR*s, comprising three *CNL*s and two *NLR*s, showed specific upregulation in the epidermis. Noteworthy is the significant downregulation of three genes encoding putative TaRNLs, carrying an RPW8-like CC domain with a proposed role as helper NLRs in immune signal transduction, upon *Fusarium* inoculation in the procambium, phloem, or outer sheath (Figure 5A and Supplementary Table 2). This suggests limited ETI activation upon non-adapted pathogen *Foc* infection, primarily in the epidermis and phloem. However, the epidermis-specific *NLR*s were upregulated in the parenchyma at 3 DPI of adapted pathogen *Fgr* infection, with L2-specific *NLR*s showing more extensive upregulation across most cell types than during *Foc* infection (Figure 5A). In addition, the widespread upregulation of *NLR*s coincides with the expression pattern of *PRR*s at 3 DPI during *Fgr* infection. Therefore, the limited upregulation of ETI in wheat’s nonhost resistance was boosted to more cell types during the susceptible interaction with *Fgr* at 3 DPI.

Systemic Acquired Resistance (SAR), induced by local PTI and ETI responses, can propagate immunity to distal tissues. Salicylic acid (SA), a key signaling molecule for SAR, is synthesized from chorismate via two independent pathways involving isochorismate synthase (ICS) and phenylalanine ammonia-lyase (PAL) (Peng et al., 2021). While single-copy *ICS* genes have been identified in rice and barley (Pál et al., 2014; Hao et al., 2018), wheat harbors three homologs sharing over 80% identity with them. However, no significant changes in the expression of wheat *ICS* genes were detected upon *Fusarium* infection (Figure 5A and Supplementary Table 2). Genes in the alternative SA biosynthesis pathway, comprising five chorismate synthases, two chorismate mutases, and numerous PALs, exhibited significant upregulation in the phloem and outer sheath upon non-adapted pathogen *Foc* infection. However, in coleoptiles infected by the adapted pathogen *Fgr*, the upregulation of these genes was delayed in the phloem and outer sheath, and these genes were substantially upregulated across most cell types at 3 DPI (Figure 5A). The biosynthesis of N-hydroxypipecolic acid (NHP), another potent signaling molecule for SAR activation (Hartmann et al., 2018), involves 1 FMO-like and 3 ALD-like genes, which were significantly upregulated in L2-derived cell types upon *Fusarium* infection, especially in the outer sheath (Figure 5A and Supplementary Table 2). Compared to *Foc*, the infection by the adapted pathogen *Fgr* induced a stronger expression of NHP biosynthesis-related genes in L2-derived cell types (Figure 5A). In summary, the upregulation of SAR signaling molecule biosynthesis pathways by non-adapted pathogen *Foc* infection was mainly confined to the phloem and outer sheath. However, the adapted pathogen *Fgr* infection induced a stronger and more widespread upregulation of these pathways at a later stage.

Regarding SA receptors, such as NPRs (Ding et al., 2018; Zavaliev et al., 2020), three *NPR1* genes, two *NPR3*, and three *NPR4* genes were differentially expressed upon *Fusarium* inoculation compared to mock inoculation (Supplementary Table 2). NPR1 functions as a co-activator, while NPR3 and NPR4 act as co-repressors in the transcriptional regulation of plant defense against pathogens (Ding et al., 2018). *NPR1* expression was fairly even across most cell types, whereas *NPR3*/*NPR4* expression was more variable, with the highest levels observed in the parenchyma (Figure 5A). Following non-adapted pathogen *Foc* infection at 2 DPI, two *NPR1* genes were significantly upregulated exclusively in chlorenchyma cells (Supplementary Table 2). Conversely, upon *Fgr* infection at 2 DPI, three *NPR1* genes were significantly upregulated in chlorenchyma, with one also showing upregulation in the epidermis, one in the inner sheath, and another in the procambium (Supplementary Table 2). In contrast to *NPR1*, four *NPR3*/*NPR4* genes were significantly downregulated in the outer sheath cells at 2 DPI (Supplementary Table 2). Moreover, at 3 DPI of *Fgr* infection, the upregulation of an *NPR1* gene and the downregulation of an *NPR3* gene occurred in the same cell type, the parenchyma. These findings indicate that adapted pathogen *Fgr* infection significantly enhanced the perception of SAR signaling in more cell types during 2 to 3 DPI, through the upregulation of activators and downregulation of repressors.

SAR activation leads to the accumulation of pathogenesis-related (PR) proteins, which are classical markers of SAR and exhibit potential *in vitro* antimicrobial activities. The expression of *PR1* and *PR4/5* genes increased mainly in the outer sheath, whereas *PR10* genes were predominantly upregulated in the phloem (Figure 5A). PR4 is a subfamily of chitinases, while PR5 are thaumatin-like proteins with putative membrane disruption function. PR10 proteins, also known as Bet v1-like proteins, have been identified as participating in the production of alkaloids and phenolics including flavonoids, by acting as binding proteins (Morris et al., 2021). Thus, the upregulation of *PR* genes was highly heterogeneous, but there were no marked differences between adapted and non-adapted *Fusarium* infections (Figure 5A).

**Supplementary Note 4. Expression pattern analysis of defensive secondary metabolite biosynthesis under fungal treatments** Beyond the involvement with SA, metabolites from the phenylpropanoid pathway also contribute to several phytoalexin biosynthetic pathways, including those for isoflavones and hydroxycinnamic acid amides (HCAAs). Isoflavones, a class of phenolic compounds, mediate important interactions with plant-associated microbes (Dixon et al., 2002; Mathesius, 2018). A novel biosynthetic pathway for isoflavones, distinct from that found in legumes, was recently identified in wheat (Polturak et al., 2023). The genes responsible for isoflavone biosynthesis, specifically *TaCHS1*, *TaOMT8*, *TaOMT3*, *TaCYP71C164*, *TaCYP71F53*, and *TaOMT6*, were notably upregulated in the outer sheath following infection by *Fusarium* pathogens (Figure 5A). In contrast, the biosynthesis of a precursor, requiring multiple *PAL*s, two *C4H*s, and three *4CL*s within the phenylpropanoid pathway, was primarily induced in the phloem (Figure 5A). Thus, the upregulation of isoflavone biosynthesis was partitioned within two spatially isolated cell types.

HCAAs, known for their antimicrobial properties and role in cell wall reinforcement, are synthesized through agmatine coumaroyl transferase (ACT), a rate-limiting enzyme (Liu et al., 2022). Two genes encoding ACT were specifically upregulated in the outer sheath following infection by the non-adapted pathogen *Foc*, whereas their expression remained unchanged upon infection by the adapted pathogen *Fgr* (Figure 5A and Supplementary Table 2). A new *TaACT* gene within the Fusarium head blight QTL-2DL region of a resistant near-isogenic line confers resistance to *Fgr* through elevated HCAA production (Kage et al., 2017). Therefore, these findings suggest that *Fgr* may prevent the upregulation of *TaACT*s in the outer sheath to reduce the production of HCAA.

Benzoxazinoids (BXDs), indole-derived specialized metabolites present in several monocot crops including wheat, maize, and rye, play a role in plant immunity against herbivorous arthropods and fungal pathogens (Batyrshina et al., 2022; Stahl, 2022). The genes encoding cytochrome P450 monooxygenases, which convert indole into the core structure of BXDs (including two *Bx2*, two *Bx3*, two *Bx4*, and two *Bx5*), were constitutively expressed in the chlorenchyma (Figure 5A). In contrast, genes encoding cytosolic dioxygenases (two *Bx6*) and glucosyl-transferases (three *Bx8/9*), which convert BXDs into storage forms, were mainly expressed in the inner sheath (Figure 5A). The expression of genes in the BXD biosynthetic pathway was slightly reduced in various L2-derived cell types following *Fusarium* infection (Figure 5A and Supplementary Table 2). However, genes upstream of BXD biosynthesis, such as those encoding indole-3-glycerolphosphate synthase (three *IGPS1* and three *IGPS3*) and the indole synthase (nine *Bx1*), were significantly upregulated in the outer sheath upon infection by both *Fusarium* species (Figure 5A). In addition, in the susceptible interaction with *Fgr*, the expression of these precursor biosynthetic genes was also upregulated in the inner sheath, where BXD storage forms are biosynthesized (Figure 5A). Collectively, under mock treatments, the biosynthesis and storage of BXDs were organized in two separate cell types. However, upon *Fusarium* infection, the production of BXD precursors was enhanced in a third cell type, the outer sheath.

Lignin, a heterogeneous polymer of monolignols, strengthens the cell wall against pathogens. In nonhost resistance to *Foc*, the biosynthesis of phenolics from phenylalanine, the upstream pathway for lignin monomer production, showed significant upregulation of numerous *PAL*s, two *C4H*s, and three *4CL*s in the phloem from 1 to 3 DPI, with some genes showing transient upregulation in the outer sheath at 1 DPI (Figure 5A). The biosynthesis of lignin monomers from phenolics in the coleoptile involves several differentially expressed genes, including a *CCoAOMT*, two *F5H*s, a *COMT,* and five *CAD*s. However, except for *CAD*s, the expression of other genes was not induced in the phloem. Instead, they were significantly upregulated in the outer sheath at 1 DPI (Figure 5A). Thus, in nonhost resistance to *Foc*, the transcriptional activation of lignin monomer biosynthesis from phenylalanine was more integrated in the outer sheath than in the phloem.

Dirigent proteins are proposed to be involved in monolignol dimerization to free lignans (Hatfield and Vermerris, 2001; Kim et al., 2012). Recently, they have also been reported to be essential for lignin polymerization of the Casparian strip in roots (Gao et al., 2023). Five putative dirigent protein-encoding genes were upregulated in the wheat coleoptile infected by *Fusarium* pathogens, with their expression patterns aligning with monolignol biosynthesis in the phloem (Figure 5A). In nonhost resistance to *Foc*, the dirigent genes were rapidly upregulated in the outer sheath at 1 DPI (Figure 5A).

However, during the susceptible interaction with *Fgr*, the upregulation of these genes in the outer sheath was diminished or delayed until 3 DPI (Figure 5A). Additionally, the induction of phenolic biosynthesis from phenylalanine was also delayed until 3 DPI in the outer sheath (Figure 5A). Thus, the infection of the adapted pathogen *Fgr* impaired the upstream production and polymerization of lignin monomers in the outer sheath.

**Supplementary Note 5. Pseudotime expression analysis of parenchyma defense strategies** It has previously been reported that the directional growth of *Fgr* hyphae is stimulated by the catalytic product of secretory peroxidases in wheat (Sridhar et al., 2020; Sridhar et al., 2023). Transcriptional upregulation of these secretory peroxidases (*TaPOD237* and *TaPOD363*) was observed in branches of the parenchyma trajectory (Supplementary Figure 8). Additionally, the expression of *PR1*, a hallmark of salicylic acid (SA)-mediated defense (Conrath, 2006; Vlot et al., 2009), was specifically induced at the border of the infection site in Arabidopsis (Jacob et al., 2023). Among the 84 PR1 genes identified in the wheat genome (Liu et al., 2023), our analysis revealed that 25 of these genes, with 11 showing unique expression patterns across different trajectory branches (Supplementary Table 5). Furthermore, nine PR1 genes were significantly upregulated in the outer sheath in response to *Fusarium* infection (Supplementary Table 2). Of these PR1 genes in the coleoptile, *TaPR1-79*, *TaPR1-51*, and *TaPR1-80* were upregulated in branches of the parenchyma trajectory, but with different amplitudes (Supplementary Figure 8). Collectively, the peroxidase-mediated chemotropism of *Fgr* and the expression patterns of *PR1* genes in wheat indicate a spatial association between fungal hyphae and parenchyma cells.

Subsequently, we aimed to ascertain the defense strategies of parenchyma cells to *Fusarium* during spatiotemporal interactions by analyzing the expression dynamics of established immune genes. The thaumatin-like protein family (PR5), with approximately one hundred members identified in the wheat genome (Sharma et al., 2022; Ren et al., 2022), included 20 *TaTLP*s with distinct expression patterns across different trajectory branches (Supplementary Table 5), 14 of which were differentially expressed in the outer sheath following *Fusarium* infection (Supplementary Table 2). Representative TLP-encoding genes, including *TaTLP14-B5*, *TaTLP16-A*, and *TaTLP32-A*, demonstrated a gradual increase in expression in branches 2 and 3, with a transient spike in branch 1 (Supplementary Figure 8). Furthermore, *TaUGT12887*, a glycosyltransferase gene conferring limited resistance to the *Fgr* mycotoxin (Schweiger et al., 2013), exhibited a gradual increase in expression in branch 2 under *Fusarium* treatment (Supplementary Figure 8). *TaSAM*, a methyltransferase gene known for contributing to FHB resistance in wheat (Malla et al., 2021), and *TaCYP71F53*, encoding a specialized wheat-specific isoflavone synthase (Polturak et al., 2023), both exhibited gradual upregulation across all branches under *Fusarium* treatment (Supplementary Figure 8). Notably, the immune response was most pronounced in branch 2, followed by branch 3, whereas the response in branch 1 was eventually downregulated after a transient upregulation. Thus, the parenchyma cells mounted immune responses of varying amplitude during common and *Fgr*-specific strategies.

However, certain genes exhibited treatment-specific upregulation patterns. *TaACT*, involved in the synthesis of HCAAs (Kage et al., 2017), was upregulated in branch 2 only under *Foc* treatment, not *Fgr* treatment (Supplementary Figure 8). *TaFROG*, a *Pooideae*-specific orphan protein that enhances resistance to *Fgr* (Perochon et al., 2015), exhibited late upregulation in branch 2 specifically under *Fgr* treatment, not *Foc* treatment. The regulation of defense genes *TaACT* and *TaFROG* appears to be opposing in infections caused by *Fgr* and *Foc*, indicating their potential involvement in susceptible processes. Furthermore, several genes encoding uncharacterized proteins relevant to disease demonstrated divergent expression patterns between *Fgr* and *Foc* treatments across each trajectory branch. An uncharacterized glycosyltransferase gene (*TraesCS2B02G040500*) displayed gradual upregulation in branch 2 and late upregulation in branch 3 mainly under *Foc* treatment (Supplementary Figure 8). The gene encoding an auxin-responsive protein with a PB1 domain (*TraesCS1D02G119500*) was upregulated late in branch 2 exclusively in response to *Fgr* treatment (Supplementary Figure 8). A gene encoding a flavin-containing monooxygenase (*TraesCSU02G099800*), potentially involved in NHP biosynthesis, showed gradual upregulation under *Fgr* treatment, particularly in branch 2 (Supplementary Figure 8). In summary, the treatment-specific upregulation of defense genes reveals the competition between immune and susceptible processes in the parenchyma defense strategies.

## Supplementary Materials

**Supplementary Table 1. Marker genes for cell annotation**

**Supplementary Table 2. Differentially expressed genes of each cell type**

**Supplementary Table 3. Enrichment analyses of differentially expressed genes**

**Supplementary Table 4. A curated list of immunity-related genes**

**Supplementary Table 5. Genes associated with parenchyma trajectory**

**Supplementary Table 6. Genes associated with chlorenchyma trajectory**

**Supplementary Table 7. Genes encoding SEID proteins in the wheat genome**

